# Dynamic observation of ^2^H labeled compounds in the human brain with ^1^H versus ^2^H magnetic resonance spectroscopy at 9.4T

**DOI:** 10.1101/2022.01.24.477582

**Authors:** Loreen Ruhm, Theresia Ziegs, Andrew Martin Wright, Claudius Sebastian Mathy, Saipavitra Murali-Manohar, Johanna Dorst, Nikolai Avdievich, Anke Henning

**Author notes:** Corresponding author:* Loreen Ruhm, High-Field MR Center, Max Planck Institute for Biological Cybernetics, Tübingen, Germany. contributed equally to this work (Loreen Ruhm, Theresia Ziegs).

## Abstract

The metabolic pathway of [6,6’-^2^H_2_]-labeled glucose was investigated with two different techniques. The first technique used direct detection of deuterium applying Deuterium Metabolic Imaging (DMI). The second technique used the indirect detection of deuterium with proton MR spectroscopy (MRS) called Quantitative Exchanged-label Turnover (QELT) MRS. For the first time, time-resolved data was acquired for both techniques in the same healthy human subjects and directly compared. The time-curves were used in a kinetic model to estimate rates of the metabolic pathway of glucose. Two different kinetic models were compared. One included only DMI data, the second one combined DMI and QELT. For the first model, a tricarboxylic acid (TCA) cycle rate of 0.69 ± 0.10 μmol·min^-1^·g^-1^ was determined. For the second model, the estimated TCA cycle rate was 0.68 ± 0.12 μmol·min^-1^·g^-1^. In addition, the rate of glutamine synthesis from glutamate could be estimated with model 2 (0.51 ± 0.15 μmol·min^-1^·g^-1^). The sensitivity of both methods was evaluated and compared to alternative techniques.

## Introduction

Altered fluxes through the metabolic pathways of glucose are of major interest in different pathologies, e.g. brain tumors. Recent publications from De Feyter et al. presented a magnetic resonance imaging (MRI) technique that can track the metabolism of deuterated glucose called DMI (Deuterium Metabolic Imaging).^1, 2^ This technique enables the non-invasive investigation of the glucose metabolic pathways by observation of ^2^H-label incorporation into downstream metabolites after oral administration of ^2^H-labeled glucose. De Feyter et al. presented data acquired from healthy subjects as well as patients with brain tumor.^1^ The alteration of glucose metabolism was clearly detectable in the tumor tissue. A challenge for the widespread application of DMI is the need for dedicated RF coils, RF amplifiers, and receive channels to detect ^2^H directly. Therefore, Rich et al. presented an alternative method called QELT (quantitative exchanged-label turnover MRS)^3^. With this method an indirect detection of the ^2^H labeling is possible. The exchange of hydrogen nuclei by deuterium nuclei in metabolites after the administration of [6,6’-^2^H_2_]-labeled glucose can be detected indirectly using proton MRS. Therefore, this technique requires proton RF coils, amplifiers, and receive channels only, which are widely available. In their publication, Rich et al. compared both methods in animal models and showed their consistency.^3^ With both techniques slightly different metabolites are detectable as detailed below.

The DMI detectable metabolites are shown in Fig. 1. This figure shows the metabolism of deuterium labeled glucose and was adopted from Lu et al.^4^ and De Feyter et al.^1^. The deuterium labeling is indicated with D (red). The [6,6’-^2^H_2_]-labeled glucose (Glc6) is converted by the glycolysis to pyruvate, which quickly exchanges with a lactate pool. DMI detectable is the ^2^H labeling of lactate in the third position (Lac3). Pyruvate can be further processed to acetyl-CoA, which enters the tricarboxylic acid (TCA) cycle and is herein converted to citrate and α-ketoglutarate. The latter can be converted to glutamate, which is labeled at the fourth position (Glu4). Glutamate can be transformed to glutamine. The ^2^H labeling at the fourth position of the combined pool (Glx4) of glutamate and glutamine (Gln4) is DMI detectable. The TCA cycle also releases deuterated water, which is also DMI detectable. To summarize, with DMI, the detectable metabolites are: deuterated water, glucose, a combined resonance consistent of glutamate and glutamine called Glx and lactate. Lactate is prominent in cancer tissue as shown by De Feyter et al.^1^, but overlaps with natural abundant ^2^H nuclei in lipid signals^5^.

**Fig. 1:**
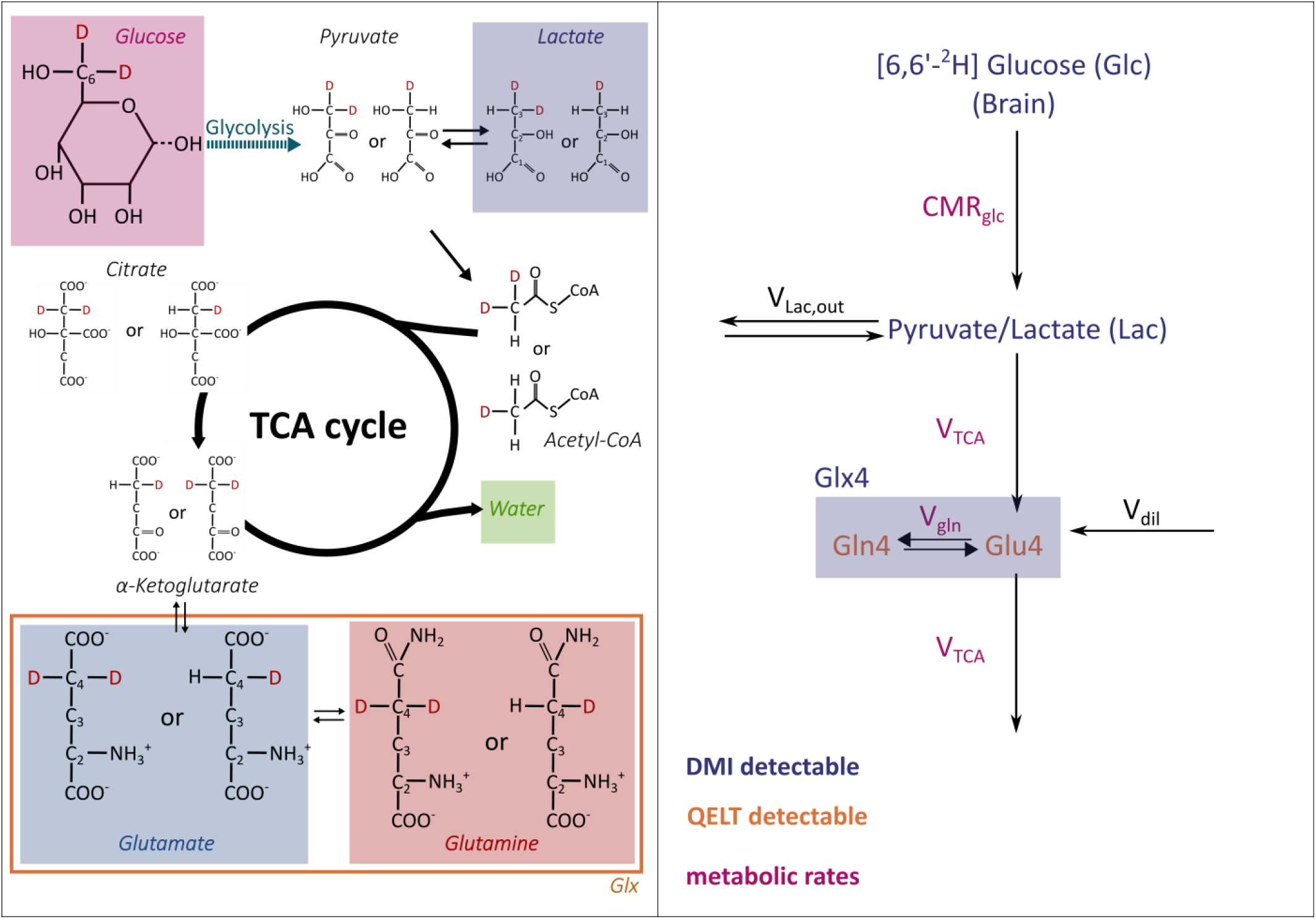
Metabolic pathway and model of [6,6’-^2^H_2_]-labeled glucose. The left side shows the metabolic pathway of [6,6’-^2^H_2_]-glucose. The MRS detectable metabolites in shortened notation are indicated with different colors. The glucose is metabolized in the glycolysis to pyruvate, which can either be processed to lactate or acetyl-CoA. Acetyl-CoA enters the TCA cycle and is herein metabolized to water and glutamate. Glutamate can be transformed to glutamine. The right side shows the metabolic models. Two different models were used to estimate the metabolic rates of metabolism of [6,6’-^2^H_2_]-labeled glucose in the human brain. The models are adapted from Lu et al.^4^, Mathy et al.^25^, and Mason et al.^26^. The first model includes only the changes in concentration of Glc6, Lac3, and Glx4 measured with DMI. The second model uses the additional information of separated Gln4 and Glu4 from the QELT measurement.

With QELT, Rich et al. detected the ^2^H uptake of glutamate and glutamine separately as well as GABA for animal experiments.^3^

In this work, we present first experiments comparing time-resolved QELT and DMI in healthy human brain. All measurements were performed at an ultra-high field strength of 9.4 T. The volunteers were measured twice with an identical oral administration of ^2^H-labeled glucose to compare both techniques. Two kinetic models are proposed to estimate metabolic rates measured from metabolizing of [6,6’-^2^H_2_]-labeled glucose in the human brain after oral administration. In the first part of the manuscript, the experimental setup as well as measurement parameters and the data post-processing is described. Afterwards, the kinetic models are explained and metabolic turnover rates are calculated. The sensitivity of both techniques was compared. The results were also compared to alternative methods as e.g. ^13^C MRS.

## Material and Methods

All measurements were performed using a 9.4T Magnetom whole-body MR system (Siemens Healthineers, Erlangen, Germany). Six different volunteers participated in this study (female: 3, average age: 28.7, max. age: 30, min. age: 26). All in vivo experiments were approved by a local ethic committee and performed after written informed consent. The following sections describe the study protocols for the corresponding measurement days. All volunteers were scanned on two separate days. The volunteers were asked to fast 9h before the start of the measurement. On the first day, the DMI experiment was performed and on the second day, the QELT single-voxel spectroscopy (SVS) data was acquired. Each volunteer orally consumed 0.75 g/kg body weight of [6,6’-^2^H_2_]-labeled glucose dissolved in 150 to 200 ml water. A blood sugar measurement (Accu-Chek *Aviva*) was performed with each volunteer prior and after the measurement. No additional measurement of blood glucose was performed.

### DMI

The DMI experiments were performed with a dual-tuned phased array coil (10 TxRx proton channels, 8TxRx/2Rx channels for deuterium^5, 6^). ^2^H is not a standard nucleus for Siemens whole body MR imaging systems. Hence the deuterium frequency needs to be externally generated and provided to the system. A frequency of 59.673 MHz and an amplitude of 8 dBm was generated by a signal generator (EXG Analog Signal Generator, NS171B, 9 kHz – 3 GHz, Agilent) and fed into the transmitter and reception boards of the scanner via a respective custom made power splitter.^5^

Before the DMI measurement, the vendor implemented, image-based 2^nd^ order B_0_ shimming was applied. A fast Ernst angle 3D magnetic resonance spectroscopic imaging (MRSI) sequence was used for deuterium metabolic imaging applying the following parameters^5^: field-of-view FoV (180×200×180) mm^3^, grid size (12×13×14), repetition time T_R_ = 155 ms, 11 averages, Hanning weighting, rectangular pulse with T_p_ = 0.5 ms, flip angle = 51 deg, vector size 512, 20 preparation scans, acquisition bandwidth 5000 Hz. The nominal voxel size is 2.94 ml. The temporal resolution is 10 min covering the whole brain (Fig. 2A). For anatomical imaging, a 3D MP2RAGE^7^ was acquired: spatial resolution (1 mm)^3^, flip angle = 4 and 7 deg, echo time TE = 2.27 ms, inversion time TI = 750/2100 ms, T_R_ = 6 ms, volume T_R_ = 5.5 s. The MP2RAGE images were corrected for B _1_^+^ inhomogeneities using an actual flip angle imaging (AFI): TR_1_ = 20 ms, TR _2_= 100 ms, TE = 4 ms and flip angle 60 deg.

**Fig. 2:**
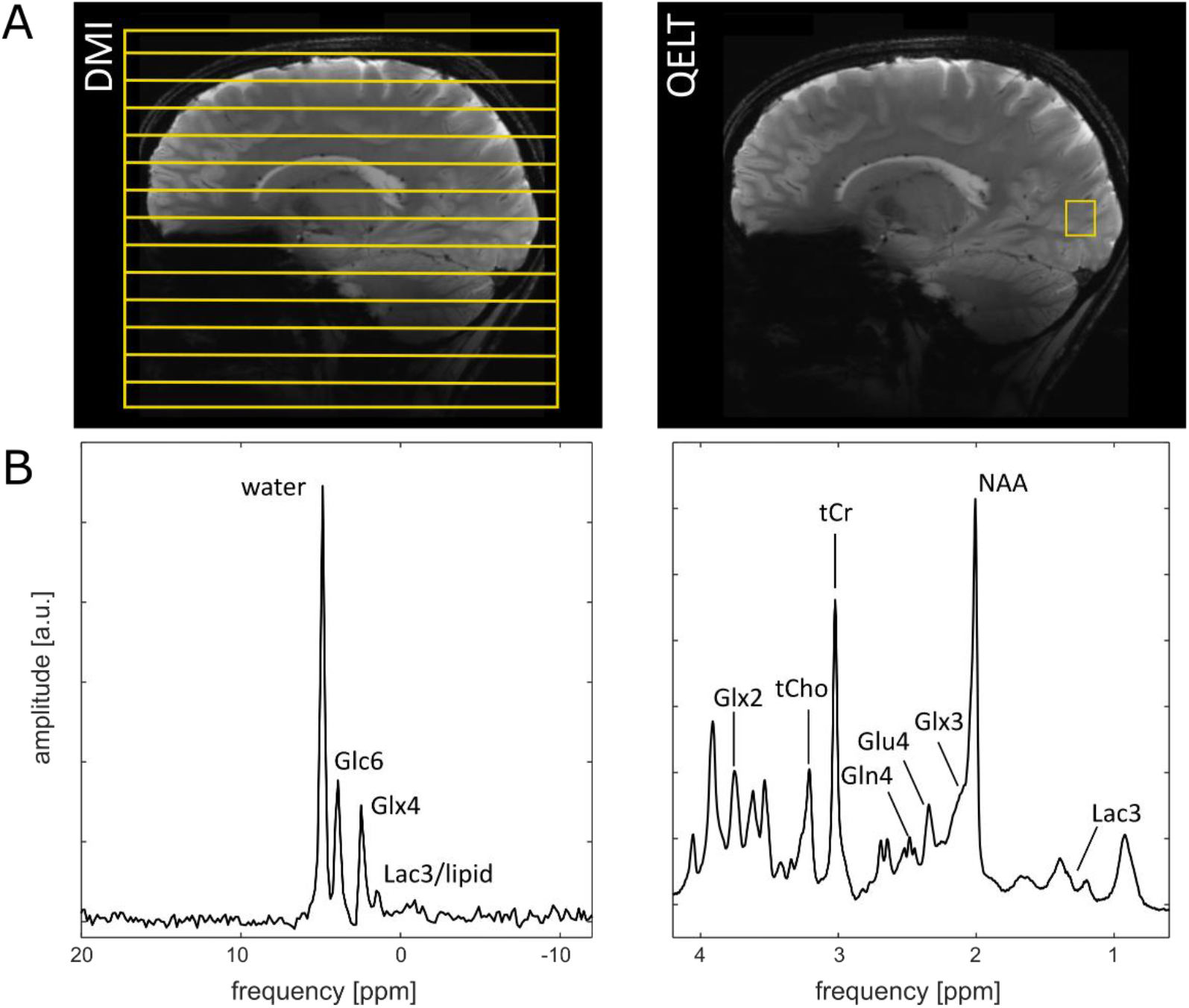
Positioning and spectral quality. The first row (A) shows the positioning of the DMI grid and QELT voxel. The second row (B) shows representative spectra for both measurements. The applied post-processing is described in the main text. Both spectra were acquired from the same volunteer.

Before the administration of ^2^H-labeled glucose, a reference DMI dataset as well as all anatomical measurements were acquired. After the administration of the glucose, multiple DMI measurements were performed each with a duration of 10 min until a total scan time of 120 min was reached. The ^2^H-labeled glucose was consumed in a supine position.

Post-processing of the DMI data was performed with in-house-written code (Matlab R2018a) and included the following steps: 3D Fast Fourier Transformation (FFT), WSVD (whitened singular value decomposition) coil combination^8, 9^, temporal zero-filling to four times the resolution, 40 Hz Gaussian time domain filter and zero-order phase correction, performed by maximizing the integral of the real part of the deuterated water resonance in the spectral domain. For signal amplitude quantification, a self-implemented version of the AMARES algorithm^10^ (Matlab R2018a) was used with a Lorentzian line shape. The first-order phase for the fit was fixed to 0.66 ms, which is determined by the sequence parameters. One zero-order phase for all resonances was fitted per spectrum. Four resonances were fitted: deuterated water, glucose (Glc6), Glx4 (= combined pool of glutamine and glutamate labeled on the fourth group, see Fig. 1) and lipids/lactate (Lac3).

The tissue content of each DMI voxel was calculated using the anatomical MP2RAGE measurement. The SPM12 algorithm^11^ was used for tissue segmentation. Afterwards, the tissue content per DMI voxel was calculated using an in-house-implemented Python script (Python 2.7).

### QELT

All QELT single voxel proton magnetic resonance spectroscopy (SVS) measurements were performed using a half-volume 3Tx/8Rx proton coil.^12^ Prior to the QELT SVS acquisition, gradient echo (GRE) scout images were acquired in sagittal and transversal direction for voxel placing. The voxel was positioned in the occipital lope (Fig. 2). FAST(EST)MAP^13^ (Fast, Automatic Shim Technique using Echo-planar readout for Mapping Along Projections) was used for first- and second-order B_0_ shimming; followed by a voxel-based power calibration.^14, 15^ The QELT SVS data was acquired using a short TE ^1^H MC-semiLASER (metabolite cycling semi Localization by Adiabatic SElective Refocusing) sequence applying the following parameters^16^: TE = 24 ms, T_R_ = 5 s, 64 averages, 512 ms acquisition time and 8 kHz bandwidth. The voxel size is (15×18×20) mm^3^ (5.4 ml). The temporal resolution of this measurement was 5:40 min. This measurement is referenced herein as metabolite spectrum. In addition, a water-reference measurement (16 averages) as well as a macromolecular (MM) measurement (TR = 8 s, 32 averages) were acquired in the exact same position. All of these measurements were performed once before the oral administration of ^2^H-labeled glucose. After the administration of glucose, multiple QELT SVS measurements as well as an additional water reference and anatomical images were acquired until a total measurement time of 120 min was reached.

The post-processing of the QELT SVS data was also performed in Matlab (R2018a) and is identical to former publications^17-19^ (for a detailed description see e.g. Murali-Manohar et al.^19^). For quantification of the signal amplitudes of the proton resonances, LCModel (V6.3-1L)^20^ was used with the following resonances in the simulated basis set (VeSPA, version 0.9.5 https://scion.duhs.duke.edu/vespa/^21^): macromolecular baseline, N-acetylaspartic acid (NAA), aspartate (Asp), phosphocreatine (PCr), creatine (Cr), glucose (Glc), glutamine-4 (Gln4), Glx2, Glx3, glutamate-4 (Glu4), glutathione (GSH), lactate (Lac), myo-inositol (mI), total choline (tCho), phosphorylethanolamine (PE) and taurine (Tau). Note that for separated groups were fitted for glutamate and glutamine: Glx2, Glx3, Gln4, and Glu4.

To calculate the tissue content of the QELT voxel, the GRE scout images were co-registered to the MP2RAGE measurement from the DMI session using the manual co-registration of SPM12^11^. Afterwards, the in-house-implemented Python script was used to calculate the tissue content per QELT voxel. The voxel placement was corrected manually if the co-registration of the images failed due to the small volume covered by the GRE images.

### Difference spectra

Difference spectra were calculated after correcting the first and second order phase and frequency alignment. The spectra were normalized by the corresponding water reference measurement for both methods. The difference spectra were calculated between the first measurement after the administration of labeled glucose and all the following measurements to avoid artifacts due to motion. The difference spectra were averaged over all volunteers (n = 6).

### Estimation of metabolite concentrations and metabolic rates

To be able to estimate metabolic rates and to make both methods comparable, two steps needed to be considered:

- Calculation of tissue concentrations of brain metabolites
- Matching of position and tissue fractions

To estimate metabolite concentrations from the DMI measurement, the measurement of natural abundant water acquired before the administration of glucose was used as a reference. In addition, saturation effects due to *T*_*1*_ *relaxation* were corrected using the T_1_ relaxation times for the deuterium labeled metabolites measured by De Feyter et al.^1^ under the assumption of field-independent T_1_ relaxation for deuterium^1, 5^. The water T_1_ relaxation time measured at 9.4 T was taken from an earlier publication^5^. *T*_*2*_ *relaxation* do not need to be considered for the DMI measurement due to the short acquisition delay of the ^2^H MRSI sequence. The equation used for the estimation of the concentrations for DMI is given in the supplemental material A1. The following label losses taken from de Graaf et al.^22^ were considered to account for the loss of ^2^H labels during the metabolic pathway: 15.7 ± 2.6 % (lactate), 37.9 ± 1.1 % (glutamate), and 41.5 ± 5.2 % (glutamine).

The estimation of the concentration of the metabolites measured with QELT SVS followed the protocol described in Murali-Manohr et al.^19^. The corresponding equation can be found in the supplement material as well (A2). The concentrations were calculated using the internal water reference of the MC-semiLASER sequence. A T_2_ correction was applied using the T_2_ relaxation times for Gln and Glu measured at the human brain at 9.4 T by Murali-Manohar et al.^19^ The T_2_ relaxation time of Glc was taken from Xin et al.^23^ measured at rat brain at a field strength of 9.4 T. No T_2_ correction was applied for Lac as no values at 9.4 T could be found in the current literature. T_1_ weighting was corrected based on the T_1_ relaxation times measured at 9.4 T by Wright et al.^24^. No T_1_ correction was applied for Lac and Glc. For these metabolites no values are available for the human brain measured at 9.4 T. Due to the relatively long TR of the 5 s, T_1_ saturation effects are expected to be small for all metabolites. The relaxation times for water at 9.4 T were taken from Hagberg et al.^7^ 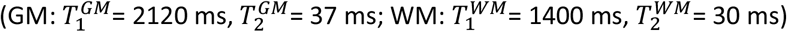.

As described above, the anatomical measurements were used to calculate the tissue fraction for each measured voxel for both methods. To calculate metabolic rates, DMI voxels with matching position and tissue content as the QELT voxel were selected. The changes in Glx4 concentration measured with both methods were compared using the averaged uptakes in Glx4 labeling over all volunteers (n=6). To be able to compare the concentrations measured at different time points between the volunteers and the two different measurement techniques, the amplitudes were interpolated to equivalent time points for each volunteer separated by 5 min for QELT and 10 min for DMI. For the comparison, the measured amplitudes of Glu4 and Gln4 of the QELT SVS were added to get the concentration of Glx4. A Pearson’s correlation analysis between the concentration changes was performed and a Bland-Altman plot calculated.

The applied kinetic models are depicted on the right side of Fig. 1. Two different models were compared. The model is adapted from Lu et al.^4^, Mathy et al.^25^ and Mason et al..^26^ The CWave software by Mason^27^ was used for modeling. The first model only used the concentration changes measured from DMI (blue). Glu4 and Gln4 are modeled as a combined pool (Glx4). CWave requires concentrations as well as fractional enrichments (FE). Therefore, the baseline concentration before the oral administration for Glx4, Gln4, Glu4, and Glc were estimated from the QELT acquisition. In the first model, the pool of natural abundant deuterium as well as the deuterated glucose were modeled as CWave drivers. Lac3 and Glx4 were modeled as CWave target concentrations. In the second model, the information from DMI and QELT was combined. Gln4 and Glu4 from the QELT data (orange) were included as additional target concentrations. The further metabolism to Glu3/Gln3 and Glu2/Gln2 was neglected due to the high ^2^H label loss as shown by de Graaf et al.^22^ The following values taken from literature were used as starting values or fixed values in the model^26^: CMR_glc_ = V_TCA_/2, V_TCA_ = 0.729 μmol · g^-1^ · min^-1^, and V_gln_ = 0.466 μmol · g^-1^ · min^-1^. The remaining dilution rates (V_Lac,out_ and V_dil_) were set to relative low rates as starting values. For model 1, V_gln_ was set to a fixed value of 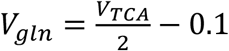. ^28^

## Results

The spectral quality of both measurements (DMI and QELT) is presented exemplary for one volunteer in Fig. 2. Shown is the spectrum from one voxel for both methods. The upper row in Fig. 2 shows the positioning of the DMI slices and the QELT voxel. Fig. 3 shows the matched position of the DMI voxel for one exemplary volunteer as well as the corresponding underlying ^2^H-Glx4 images in sagittal and transversal direction. The Glx4 images are normalized by the natural abundant water signal measured for each voxel before the oral administration of deuterated glucose. The tissue content of grey matter (GM), white matter (WM), and cerebrospinal fluid (CSF) per QELT voxel and matched DMI voxels are listed in Tab. S1 of the appendix. The DMI tissue content was corrected for the point-spread-function (PSF) of the MRSI measurement. An exemplary LCModel fit as well AMARES fit for QELT and DMI can be found in Fig. 4.

**Fig. 3:**
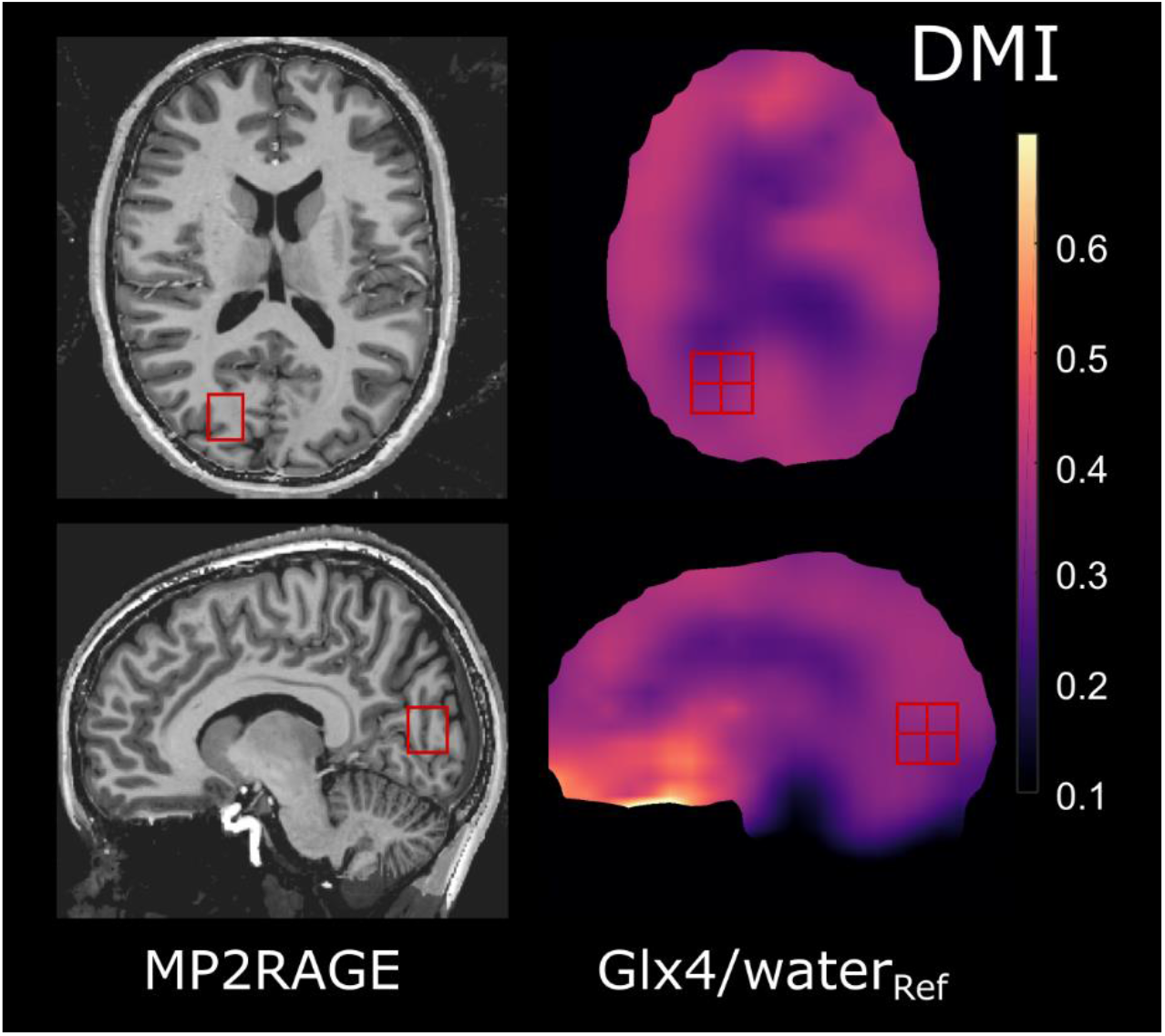
Position of matched DMI voxel. The left side shows the QELT voxel positioning for an exemplary volunteer. On the right, the corresponding DMI images are shown with the positioning of the matched DMI voxel. The ^2^H images were acquired 80 min after the oral intake of ^2^H labeled glucose and normalized to the natural abundant water signal *water*_*ref*_acquired before the oral intake of deuterated glucose.

**Fig. 4:**
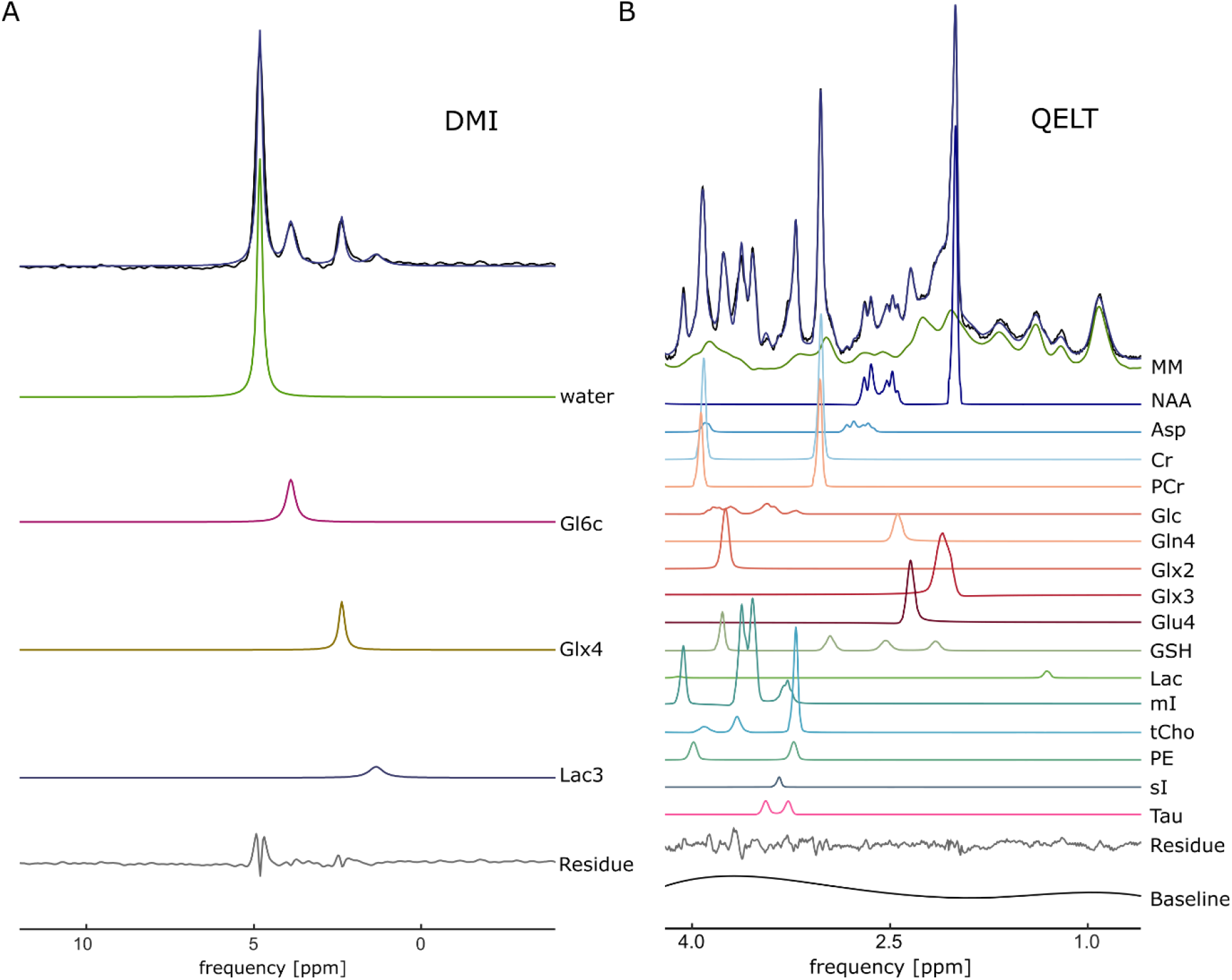
AMARES and LCModel fit of the DMI and QELT data. The left side (A) shows the AMARES fit for an exemplary DMI measurement (averaged over eight voxels). The right side (B) shows the LCModel fit of the QELT data of the same volunteers. Both spectra were acquired approx. 80 min after the oral administration of [6,6’-^2^H_2_]-glucose. All fitted metabolites per method are listed in the figure.

As described in the introduction, with DMI, water, glucose (Glc6), as well as a combined resonance for glutamate (Glu4) and glutamine (Gln4) labeled as Glx4 are separately detectable. The Glx resonance stems from the fourth group in Gln and Glu. The lactate resonance overlaps with signals arising from lipids surrounding the head and should only show a weak signal in healthy human tissue.^1^ Lactate is labeled on the third position (Lac3). The QELT spectra is shown on the right side. In an earlier publication from Rich et al.^3^, time curves of Glu4, Gln4, combined Glx, GABA (*gamma*-aminobutyric acid) and NAA could be presented for *in vivo* measurements at a rat brain. In accordance with the metabolic pathways shown in Fig. 1, Glu4, Gln4, and Glx4 should show a gradual enrichment after the administration of [6,6’-^2^H_2_]-labeled glucose.

The labeling uptake curves measured in this study are shown in Fig. 5A. For QELT, amplitude-time-curves are displayed for Glu4, Gln4, Glx3 and Glx2. GABA was not detectable for the presented human brain QELT data. The individual glucose groups could not be fitted separately. All QELT amplitudes were normalized by the internal water signal of the MC-semiLASER sequence. The strongest decrease in the ^1^H spectrum due to ^2^H labeling is visible for Glu4. Potential minor labeling uptakes may be detectable for Gln4. No uptake is detectable for Glx2 and Glx3, which is in agreement with de Graaf et al.^22^

**Fig. 5:**
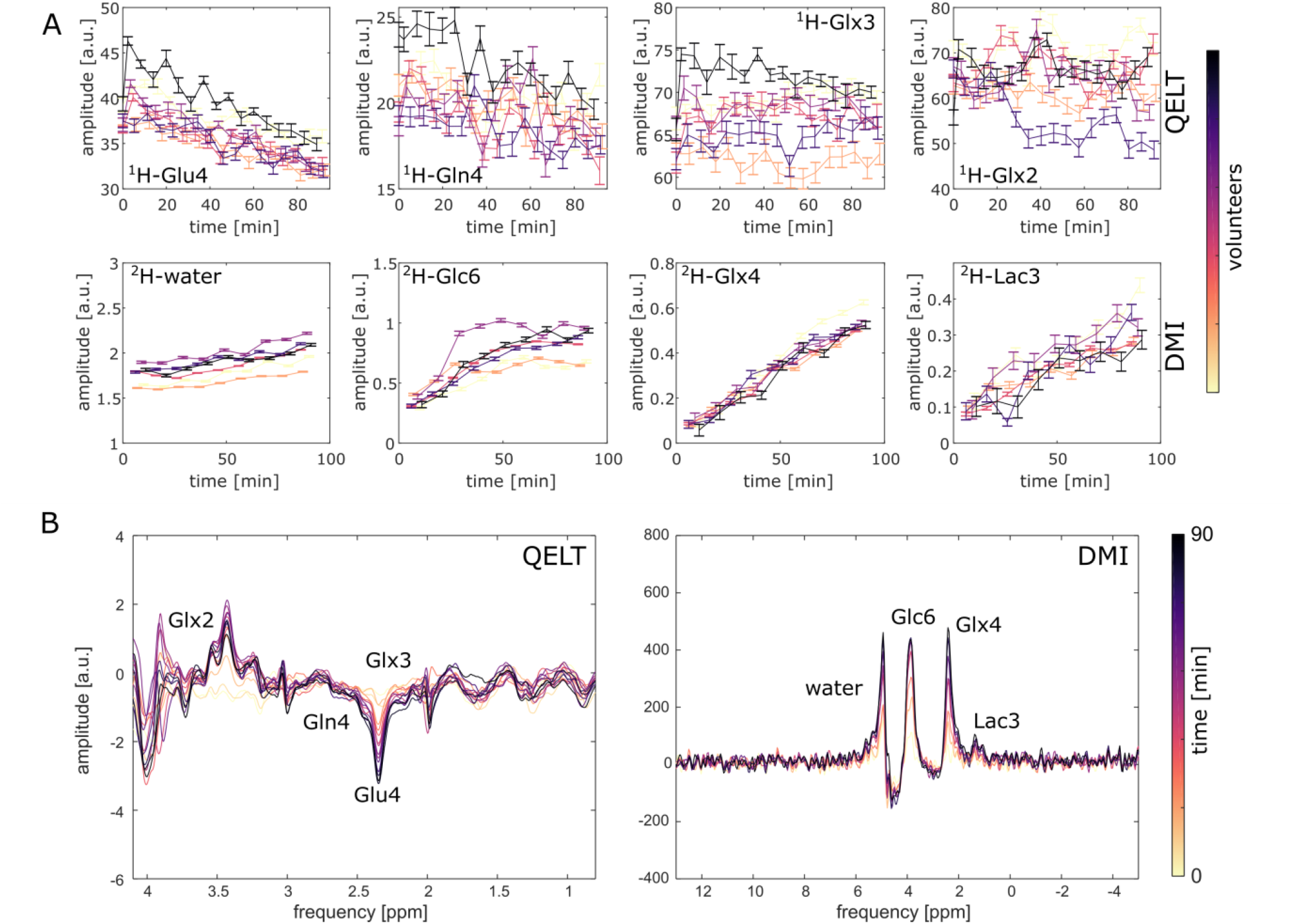
Temporal resolved QELT and DMI. The top rows (A) show the ^1^H amplitudes for Glu4, Gln4, Glx3, and Glx2 measured with QELT and the ^2^H amplitudes of deuterated water, Glc6, Glx4, and Lac3 measured with DMI of all volunteers (n = 6). The last row (B) shows the difference spectra for DMI and QELT averaged over all volunteers (n = 6).

The detectable metabolites for DMI are ^2^H-labeled water, Glc6, Glx4, and Lac3. All DMI signal amplitudes were normalized by the water signal of the reference measurement before the oral administration of ^2^H-labeled glucose to avoid RF coil loading differences, transmit B_1_^+^ or receive sensitivity B_1_^-^ weighting between the averaged DMI voxel or different volunteers. All four metabolites show a clear uptake in ^2^H-labeling. The uptake in water follows a linear trend, whereas the signal of Glx4 and Lac3 seem to saturate slowly towards long times (> 70 min) after the administration of [6,6’-^2^H_2_]-labeled glucose. Glc6 increases fast at the beginning, but reaches a maximum after approximately 70 min. Afterwards, the labeling decreases or stays relatively constant within the time frame of the study. The Glc6 uptake shows a high variance between the different volunteers, which does not seem to translate into a high variance of ^2^H Glx4 and Lac3 uptake, which are very consistent between the different volunteers.

### Difference spectra

Figure 5B shows the difference spectra calculated for the single voxel of the QELT measurements (left) averaged over all volunteers (n=6) as well as the averaged difference spectra from the matched ^2^H MRSI voxels (right). As, described earlier, for the QELT difference spectra, ^2^H labeling translates in a decrease of signal in the corresponding proton resonance. A clear consistent decrease can be seen in the region of Gln4/Glu4. No clear detectable decrease is visible in the region of Glx3 and Glx2. For the matched DMI voxels clear ^2^H uptakes are visible for water, Glc6, Glx4 and Lac3.

### Comparison of Glx4 labeling

The change in concentration of Glx4 measured with DMI was calculated as well as the concentration change due to ^2^H labeling for Gln4 and Glu4 measured with QELT SVS. Gln4 and Glu4 were summed to calculate the Glx4 uptake of the QELT measurement. As only the concentration changes due to ^2^H labeling uptake can be compared, the starting concentration of Glu4 and Gln4 measured with QELT was subtracted from the signal amplitudes. To avoid high signal variance in the baseline concentration, the starting point signal was estimated by a linear fit of the Gln4 and Glu4 signal amplitudes of the averaged concentration changes of all volunteers.

Fig. 6A shows the mean changes in ^2^H labeled Glx4 concentration measured with QELT and DMI across all volunteers. The presented errors are s.e.m. (standard error of the mean). The Pearson’s correlation analysis shows a high correlation (R = 0.91) between both measurements. The Bland-Altman plot shows a small bias between both measurements of (−0.19 ± 0.37) mM. Note again that the effect of label loss according to de Graaf et al.^22^ was corrected for the presented concentrations.

**Fig. 6:**
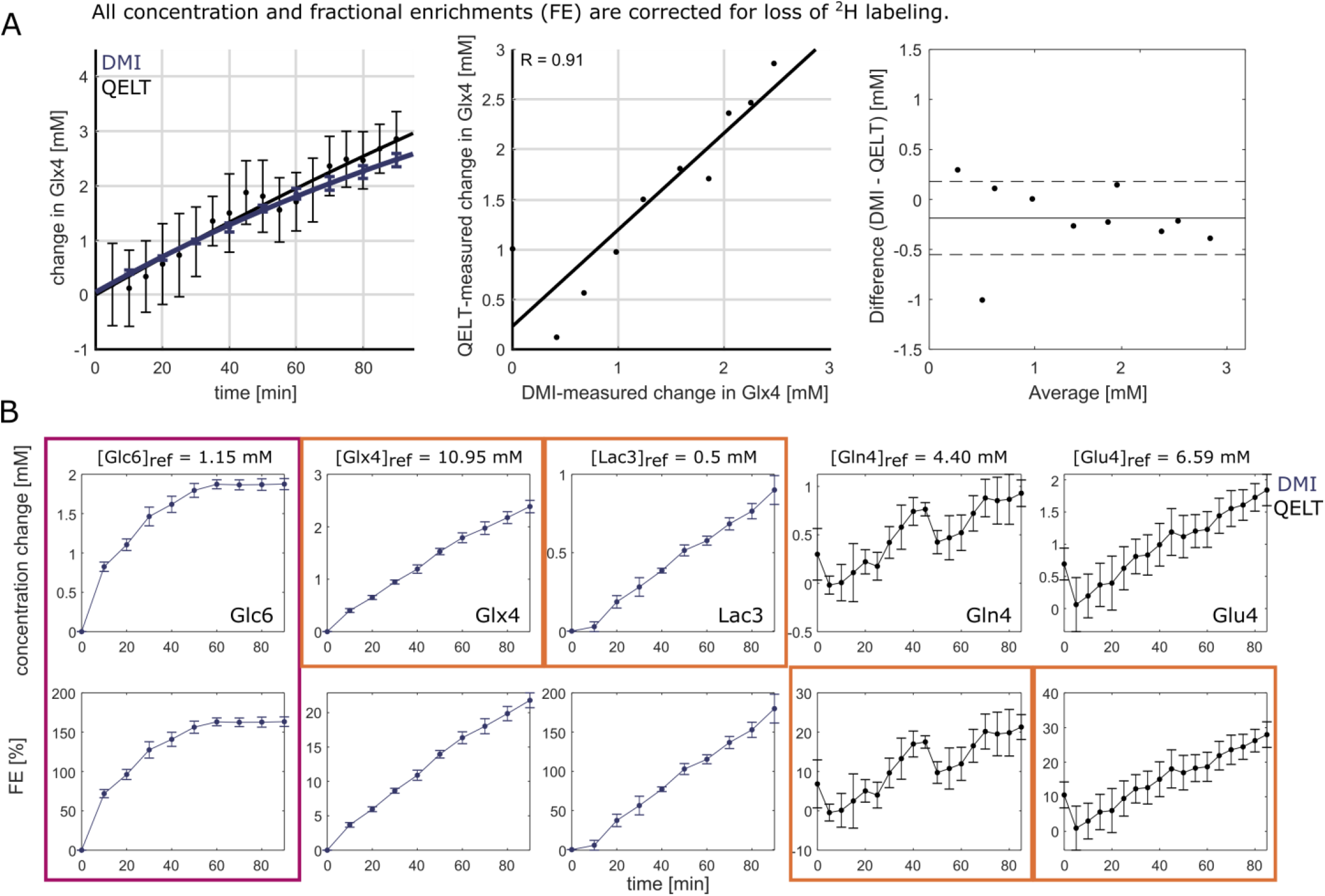
Comparison of the Glx4 concentration changes and model input concentrations and fractional enrichments. Shown are the mean changes in Glx4 due to ^2^H labeling measured with DMI and QELT averaged across all volunteers (n = 6, A). As a visual aid, a curve was fitted to both plots (*F*(*t*)=*A*_*M*−_(*A*_*M*−_*A*_0_) ·exp(−*k*· *t*) see Rich et al.^3^). A correlation plot as well as a Bland-Altman is presented to compare the Glx4 concentration changes measured with QELT and DMI. The lower rows (B) show the input concentration changes and fractional enrichment (FE) for Glc, Glx4, Lac, Glu4, and Gln4. All concentrations and fractional enrichments are corrected for ^2^H-label label loss according to de Graaf et al.^22^ The concentration changes and FE used in the model are labeled by boxes. Glu6 was used as driver data (purple) and Glx4, Lac3, Gln4, and Glu4 as target data (orange).

### Metabolic rates

The mean concentration changes across all volunteers as well as the corresponding fractional enrichments (FE) needed as input for metabolic modeling in the CWave software can be found in Fig. 6B for Glx4, Glc6 and Lac3 measured with DMI and Gln4 and Glu4 measured with QELT. To calculate fractional enrichments, the concentration for Glx, Gln, Glu, Glc, and Lac before the administration of labeled glucose were estimated from the QELT SVS data. All concentrations and FE were corrected for the expected label loss.^22^ Lac3 was corrected by the signal intensity of the resonance at t = 0 min to avoid contamination from lipids by subtracting the natural abundant signal at around 2.4 ppm. The lipid signal is not expected to change due to the glucose metabolism with in the time-frame of the study.^5^ The ^2^H concentration changes and fractional enrichment were fed into CWave using the metabolic models presented in Fig. 2. The used parameter files are given in the appendix (A3). The resulting metabolic rates are listed in Tab. 1. The error estimates were calculated using Monte Carlo simulations (n = 200). The corresponding fits of the time-curves are shown in the supplement material (Fig. S1). For the first model, the estimated value for the tricarboxylic acid (TCA) cycle rate V_TCA_ is 0.69 ± 0.10 μmol·min^-1^·g^-1^. For second model, the estimated TCA cycle rate is 0.68 ± 0.12 μmol·min^-1^·g^-1^ and the rate of synthesis glutamine from glutamate is 0.51 ± 0.15 μmol·min^-1^·g^-1^. The cycle rates of the TCA cycle agree well between both models.

**Tab. 1:** Metabolic rates for [6,6’-^2^H_2_]-labeled glucose. Listed are the estimated metabolic fluxes for both models shown in Fig. 2. The errors were estimated from Monte Carlo simulations using CWave.

| Metabolic rates [$\mu\text{mol} \cdot \text{min}^{-1} \cdot \text{g}^{-1}$] | *Model 1* | *Model 2* |
| --- | --- | --- |
| $V_{TCA}$ | $0.69 \pm 0.10$ | $0.68 \pm 0.12$ |
| $V_{gln}$ | - | $0.51 \pm 0.15$ |

## Discussion

In the presented study, a temporally resolved proton SVS based indirect detection method (QELT) for ^2^H-labeling uptake was compared to a direct ^2^H detection method (DMI) for the first time in human brain data. The measured ^2^H labelling uptakes of Glx4 proved to be consistent between both methods. With the combined information from both methods, the measured concentration could be used in kinetic models to obtain turnover rates of the metabolic pathways of glucose.

### Comparison QELT versus DMI: data quality, sensitivity and detectable metabolites

Following de Graaf et al., the relative experimental sensitivity was defined:^2^

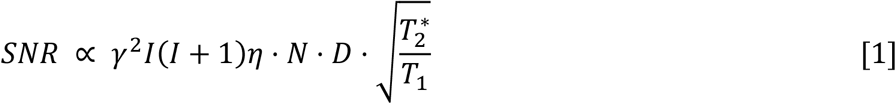

where γ is the gyromagnetic ratio in MHz/T, *I* the nuclear spin, η the potential signal enhancement using the nuclear Overhauser effect or polarization transfer, N the number of nuclei, D the decoupling efficiency and T_2_* and T1 the relaxation times.

If the experimental sensitivity of QELT and DMI are compared at 9.4 T using equation [1], the relative sensitivity DMI/QELT is 0.19 (^2^H: γ = 6.54 MHz/T, I = 1, T 2* neglected, T = 140 ms (Glx)^1^, η = 1, N = 1.5, D = 1 (see also de Graaf et al.^2^); ^1^H: γ = 42.58 MHz/T, I = ½, T_2_* neglected, T = 1300 ms (Gln/Glu)^24^, η = 1, N = 1.5, D = 1). The influence of the T_2_/T_2_* relaxation depends on the sequence. Assuming comparable short-TE ^1^H and ^2^H MRSI sequences, the effect can in both cases be neglected. Therefore, it can be concluded that QELT has a five times higher experimental sensitivity at 9.4 T. This matches with the values measured by Rich et al. in rat brain at 9.4 T.^3^ Nevertheless, our study showed that the detection of labeling changes with QELT can be challenging due to the signal overlap in the ^1^H spectrum. The separate groups of Glc were not detectable with QELT in our study neither was an uptake in Lac3.

### Fractional enrichments of glutamate and glutamine

The measured relative concentration uptake of Glx was compared between both measurements and proved to be consistent. After correcting for label losses, a labeling uptake of 21.8 ± 1.1 % was detected for Glx4 90 minutes after the oral administration of [6,6’-^2^H_2_]-glucose using DMI, and 21.3 ± 3.2 % for Gln4 and 28.0 ± 3.7 % for Glu4 measured with QELT. After 40 min, the fractional enrichments reached 10.9 ± 0.7 % (Glx4), 17.0 ± 3.3 % (Gln4), and 15.0 ± 5.0 % (Glu4). A previous study by Rich et al. that applied DMI and QELT in rats using an infusion of 1.95 g/kg body weight of [6,6’-^2^H_2_]-glucose reported 9 % enrichment for Glx, 11 % for Glu and 8 % for Gln after 45 min.^3^ No correction for labeling loss is mentioned by Rich et al. Overall the values measured in this study are slightly higher, which can potentially explained by differences between rat and human metabolism. Due to the differences in the experimental setup, a comprehensive comparison of the fractional enrichments is not possible.

As the values for the fractional enrichment presented in this study are corrected for ^2^H label loss, they can be compared to ^13^C studies. Only very few ^13^C studies use the oral administration of labeled glucose. Moreno et al. reported a fractional enrichment of Glu4 of 16 ± 2 % after the oral administration of 0.65 g/kg body weight of [1-^13^C]-glucose measured after approximately 130 min on human volunteers at 1.5 T.^30^ The value is lower compared to this study. Potentially, this can be partly explained by the lower amount of administered glucose. Mason et al. reported a fractional enrichment of approx. 18 % in humans after 150 min.^27^ Other studies that reported fractional enrichments used infusion of [1-^13^C]-glucose. These studies reported maximal enrichment of 15 % to 30 %^26, 31-35^. The value measured in this study is in a similar range.

Previous human brain ^13^C studies reported also fractional enrichments for Gln4 measured with indirect and direct ^13^C-labeled MRS. These reported values are in the range of 14 – 25 %.^26, 27, 33^ This is in agreement with the value presented in this study.

Overall, the comparison of fractional enrichments is limited by the different time-range covered, amounts of glucose, subject of interest, and experimental setups (infusion/oral administration).

### Metabolic rates

By combining the information from QELT and DMI, we were able to build a kinetic model to estimate metabolic rates after the oral administration of [6,6’-^2^H_2_]-glucose. We were able to detect the concentration changes of Glx4, Glc6, and Lac3 from DMI and concentration changes of Glu4 and Gln4 as well as the baseline concentration for Glc, Lac3, Gln4, and Glu4 before the oral administration of labeled glucose from QELT.

DMI in combination with infusion of [6,6’-^2^H_2_]-glucose was used by Mathy et al. to estimate metabolic rates in the rat brain.^25^. To the best of our knowledge, this is the only study to this point that used ^2^H-labeled MRS for the estimation of metabolic rates. They reported a rate of the TCA cycle V_TCA_ of 1.06 ± 0.20 μmol·min^-1^·g^-1^ m in the rat brain. This value is higher compared to the rate estimated in this study in the human brain.

De Graaf et al. showed that the kinetic isotope label effect (KIE) for deuterated glucose is small.^22^ The KIE describes the potential change in metabolic rates due to the replacement of atoms in the metabolites by heavier isotopes.^22^ Therefore, we can compare our findings to studies using ^13^C labeled glucose. These studies reported values for V_TCA_ of 0.73 ± 0.19 μmol·min^-1^·g^-1^ presented by Mason et al.^26^, 0.695 – 0.895 μmol·min^-1^·g^-1^ by Mason et al.^36^, 0.77 ± 0.07 μmol·min^-1^·g^-1^ by Shen et al.^37^, 0.55 ± 0.04 μmol·min^-1^·g^-1^ by Boumezbeur et al.^38^, 0.4 – 0.58 μmol·min^-1^·g^-1^ by Mason et al.^27^, and 0.80 ± 0.10 μmol·min^-1^·g^-1^ (GM) and 0.17 ± 0.01 μmol·min^-1^·g^-1^ (WM) by Mason et al.^35^ All studies beside the one by Boumezbeur et al. were performed in the human brain. Boumezbeur et al. performed their study on monkey brain. For the rate of synthesizing glutamine from glutamate V_gln_, reported rates are in the range of 0.47 μmol·min^-1^·g^-1^ (95 % confidence interval: [0.139, 3.094] μmol·min^-1^·g^-1^) reported by Mason et al.^26^, 0.33 – 0.37 μmol·min^-1^·g^-1^ by Mason et al.^36^, 0.32 ± 0.05 μmol·min^-1^·g^-1^ by Shen et al.^37^ and 0.21 – 0.27 μmol·min^-1^·g^-1^ by Mason et al.^27^ for measurements at the human brain. The estimated value for V_TCA_ of this study is in the same range as the previous publications. The rate for the conversion between glutamine and glutamate V_gln_ is slightly higher than the values reported before.

The low number of volunteers in this and the previous studies should be considered when comparing the metabolic rates.

### Limitations of the study

The replacement of protons by ^2^H nuclei were not fully considered in the VeSPA simulated spectra that was used to fit the QELT data. Instead proton only spectra have been included in the basis set for glutamate-4 (Glu4), glutamine-4 (Gln4), and their combined signals Glx2 and Glx3. The replacement of protons by deuterium nuclei at a specific position inside the molecule has two effects: (i) a signal decrease in the QELT spectra for the corresponding chemical group itself and (ii) a change in the structure of multiplets due to change in scalar coupling, which can potentially induce changes in signal intensity, phase, line width, and chemical shift of the spectral pattern of the different coupling partners. To test how the second effect changes the basis set, additional simulations were performed in VeSPA. For the assumption that the J-coupling between protons and deuterium is negligible, mainly the change in proton-proton coupling induced by replacement of protons by deuterium nuclei and hence the reduction in the number of coupled protons has to be considered. Respective simulated VeSPA basis sets for glutamate and glutamine are shown in the appendix (Fig. S2). As expected, the multiplet structure changes. Nevertheless, as the linewidths at 9.4 T are relatively high, this mainly results in broader/narrower resonances.

The basis sets in Fig. S2 still neglect the coupling between deuterium and proton. As presented by De Feyter et al., ^2^H-^1^H scalar couplings are expected to be 6.5 times smaller than ^1^H-^1^H scalar coupling due to the smaller gyromagnetic ratio, which would yield J-coupling constants below 1 Hz.^29^ Therefore, only minor to negligible effects are expected if the deuterium is included in the basis set.

Since the focus of the herein performed metabolic modelling analysis was on Glu4 and Gln4, the respective signal decrease due to the direct replacement of protons by deuterium nuclei in this group was most relevant and fully considered by the proposed proton only basis spectra for the quantitative analysis in our study. Due to substantial spectral overlap in a proton spectrum neither glutamate nor glutamine at the second and third group can be distinguished nor spectra with no, one or two ^2^H labels incorporated at the fourth group. Hence only pure proton spectra were modelled and glutamate and glutamine was jointly fitted for the second and third groups Glx2 and Glx3. Considering the different impact of ^2^H labelling on the signal intensity of Glu4, Gln4, Glx3 and Glx2 groups the most stable fitting result has been achieved by fitting these resonances individually as applied in this study.

Assumptions were also made for the used T_1_/T_2_ relaxation times for deuterium and some of the proton metabolites as mentioned in the methods part. These assumptions can lead to slight deviations in the estimated metabolite concentrations.

Another limitation of this study is the relatively short overall measurement time. Mason et al. showed that the precision of the measurement of metabolic rates can be highly improved by long measurement times of 3h.^27^ These long measurement times were not possible for human subjects within the requirements of the local ethic committee.

A potential next step would be the interleaved measurement of DMI and QELT with a double-tuned coil. At the time of this study, this was technically not possible at the 9.4 T Siemens whole-body scanner due to the need of an external signal generation for the ^2^H frequency.

### Comparison of ^2^H labeling to ^13^C and ^18^F labeling of glucose

Alternative methods to track the consumption of labeled glucose include FDG PET (fluorodeoxyzglucose positron-emissions tomography), and, as mentioned above, direct and indirect ^13^C MRS. FDG PET has the disadvantage that a radioactive tracer is required, which adds an additional burden to the patient. FDG PET can also not be used to track metabolic pathways of glucose.^39^

Therefore, the most relevant alternative method is ^13^C-labeled MRS. De Graaf et al. compared the relative sensitivity of direct ^2^H MRS and direct ^13^C MRS.^2^ They estimated that the experimental sensitivity (equation [1]) of direct ^2^H MRS is approximately twice as high as for direct ^13^C MRS assuming single-labeled ^13^C-glucose. In addition to the low experimental sensitivity, ^13^C MRS is also technically challenging as dedicated coils would be required as well as broadband proton decoupling.^40^ In addition, the broad frequency spread in ^13^C MRS resolves spectral lines very well, but leads to challenges to record localized spectra when ^13^C is directly detected.

It should be noted at this point that the experimental sensitivity of both techniques can be improved by using tracers with more labels (increases N). Nevertheless, the specificity of the method decreases with the number of labels. The labeling of different metabolic groups cannot be distinguished anymore. Therefore, we will focus on single-labeled glucose for ^13^C and double-labeled glucose for ^2^H in this discussion.

The sensitivity of ^13^C-labeling detection can also be improved by using indirect ^13^C detection with ^1^H MRS. Most of the studies that apply this technique use POCE (proton observed carbon edited).^41^ POCE uses the coupling of ^13^C nuclei to protons to detect the ^13^C labeling. The disadvantage of POCE is that it also requires a doubled tuned ^13^C/^1^H-coil as well as broadband decoupling for spectral editing.^40^ An equivalent method to QELT for the detection of ^13^C-labeling was proposed by Boumezbeur et al.^38^ This technique uses conventional proton MRS without the need for heteronuclear decoupling and without the need of a double-tuned RF coil. As QELT and the method proposed by Boumezbeur, both use conventional ^1^H MRS to detect the labeling effects, the experimental sensitivity is identical in terms of gyromagnetic ratio, spin, potential signal enhancement, decoupling, and relaxation times. Only two differences need to be minded: the effect of the labeling and labeling losses. Using [6,6’-^2^H_2_]-glucose, there are two labels that can be potentially detected with QELT compared to one label using single-labeled ^13^C-glucose. Nevertheless, the higher loss rate of the deuterium label reduces this effect. For Glu4 and Gln4, this label loss is approximately 40 % according to de Graaf et al.^22^ The ^2^H and ^13^C labeling also lead to different effects in the ^1^H spectrum: the exchange of protons for ^2^H labeling and the different change in scalar coupling for both methods. E.g., the ^13^C labeling in glutamate results into two additional peaks in the proton spectrum.^38^ A direct comparison of both methods would be potentially of interest.

## Conclusion

Significant labeling uptakes for Glx4, Gln4, Glu4, Glc6, Lac3 and deuterated water were detected with direct and indirect ^2^H-labeled MRS for six volunteers at 9.4 T after the oral administration of [6,6’-^2^H_2_]-glucose. The resulting time-curves for the labeling uptake in Glx4 were consistent between both methods. The combined information from both methods allowed the estimation of metabolic rates of the TCA cycle. The rates proved to be consistent with previous reported values.

## Acknowledgements

Funding by the ERC Starting Grant (SYNAPLAST MR, Grant Number: 679927) of the European Union and the Cancer Prevention and Research Institute of Texas (CPRIT, Grant Number: RR180056) is gratefully acknowledged. We want to thankfully acknowledge also the help of Edyta Leks with SPM and the help of Graeme Mason with CWave. Part of the data was included in an earlier publication (Ruhm et al.^5^).

## Author contribution statement

L Ruhm: Original Idea, Conceptualization, Methodology, Software, Data Curation, Formal Analysis, Writing – Original Draft; T Ziegs: Original Idea, Conceptualization, Methodology, Software, Writing – Review & Editing; AM Wright: Software; S Murali-Manohar: Software; J Dorst: Software; CS Mathys: Formal Analysis; N Avdievich: Methodology; A Henning: Original Idea, Resources, Supervision, Funding acquisition, Writing – Review & Editing

