## Supplemental Material for "Dynamic observation of ^2^H labeled compounds in the human brain with ^1^H versus ^2^H magnetic resonance spectroscopy at 9.4T"

Supplement material:

**Tab. S1:** Estimated tissue fractions for the QELT voxel and the matched DMI voxels

| Volunteer | QELT | | | DMI | | | |
|---|---|---|---|---|---|---|---|
| | GM [%] | WM [%] | CSF [%] | GM [%] | WM [%] | CSF [%] | N Voxel |
| 1 | 55 | 44 | 1 | 47 | 44 | 9 | 8 |
| 2 | 54 | 46 | 0 | 53 | 39 | 5 | 8 |
| 3 | 47 | 52 | 0 | 42 | 52 | 6 | 8 |
| 4 | 58 | 40 | 2 | 48 | 43 | 9 | 8 |
| 5 | 40 | 60 | 0 | 38 | 56 | 4 | 8 |
| 6 | 46 | 49 | 5 | 38 | 49 | 13 | 8 |

**Tab. S1:** Estimated tissue fractions for the QELT voxel and the matched DMI voxels. The tissue fractions for QELT were calculated using the MP2RAGE images and a self-implemented Python script. The tissue fractions for the QELT voxel were used together with the position of the QELT voxel to find matching DMI voxels.

**Appendix A1:** Calculating concentrations from DMI metabolites amplitudes

A concentration of 13.7 mM was assumed for the natural abundant deuterated water signal following Lu et al.[4] The following equation was applied to calculate metabolites concentrations $[M]$ from DMI metabolites amplitudes $A_M$:

$$[M] = \frac{A_M \cdot (1 - \cos\alpha \ e^{-TR/T_{1,M}})}{(1 - e^{-TR/T_{1,M}})} \cdot \frac{(1 - e^{-TR/T_{1,water}})}{A_{water} \cdot (1 - \cos\alpha \ e^{-TR/T_{1,water}})} \cdot \frac{[water] \cdot N_{water}}{N_M} \cdot \frac{1}{1 - L_M}$$

In the equation, α is the flip angle and TR the repetition time of the DMI sequence. $T_{1,M}$ and $T_{1,water}$ are the longitudinal relaxation times of the metabolite and water. The metabolites $T_1$ were taken from De Feyter et al.[1], the water $T_1$ was taken from an earlier publication of ours[5]. $N_M$ are the maximum number of deuterium nuclei per molecule: two for glucose, two for lactate and two for Glx4. In addition, label loses $L_M$ were considered for lactate, glutamate and glutamine following de Graaf et al.[22]: 15.7 ± 2.6

(lactate), 37.9 ± 1.1 (glutamate), and 41.5 ± 5.2% (glutamine). Taken the baseline concentrations measured with QELT before the oral administration of 2H-labeled glucose, an average label loss was calculated for Glx4. No label loss was assumed for deuterated glucose. The water was assumed to be dominated by single-labeled heavy water, therefore the number of deuterons per molecule $N_{water}$ equals one.

**Appendix A2:** Concentration for metabolites estimated from QELT.

The equation to estimate the concentrations for metabolites using QELT was taken from Murali-Manohar et al.[19]:

$$[M] = A_M \cdot [water] \cdot \frac{2}{1+F_s} \cdot \frac{\left(f_{GM} \cdot \alpha_{GM} \cdot R_{water,GM} + f_{WM} \cdot \alpha_{WM} \cdot R_{water,WM} + f_{CSF} \cdot \alpha_{CSF} \cdot R_{water,CSF}\right)}{(1 - f_{CSF}) \cdot R_M} \cdot \frac{1}{1 - L_M}$$

where f represents the tissue fractions within the QELT voxel (see Tab. S1) and α the relative densities of MR-visible water (GM: 78%, WM: 65%, CSF: 97%). $[water]$ is the water concentration in the human brain, which is equal to 55.126 M. $(1 - f_{CSF})$ accounts for partial-volume correction and $2/(1 + F_S)$ for the multiplication of the even numbered acquisitions with a scaling factor $F_S$. $R$ represents the relaxation correction factor:

$$R = e^{-\frac{TE}{T_2}} \cdot \left[1 - e^{-\frac{TR}{T_1}}\right]$$

$A_M$ is the signal amplitude obtained from LCModel fit divided by a preset number (40873). In addition, the concentrations were corrected for the label loses $L_M$ following de Graaf et al.[22]

**Appendix A3:** CWave parameter file

*Model 1*

```
function [comb,conc,driv,rate,initcond,targ,notes] = fun(x)
%/**** Rates ****************************************/
rate(1).V = 3.645000e-01;
rate(1).Expression = 'Vtca/2';
rate(1).iter_ans = 'n';
rate(1).name = 'CMR';
rate(1).units = 'umol/min/g';
```

```
rate(1).notes = 'Notes';
rate(1).guess = 0.000000e+00;
rate(1).delta = 1.000000e+00;
rate(1).upper_limit = 1.000000e+07;
rate(1).lower_limit = 0.000000e+00;

rate(2).V = 7.290000e-01;
rate(2).Expression = '';
rate(2).iter_ans = 'y';
rate(2).name = 'Vtca';
rate(2).units = 'umol/min/g';
rate(2).notes = 'Mason et al. (1995)';
rate(2).guess = 0.000000e+00;
rate(2).delta = 3.000000e-01;
rate(2).upper_limit = 2.000000e+00;
rate(2).lower_limit = 0.000000e+00;

rate(3).V = 7.289854e-01;
rate(3).Expression = 'Vtca-Vdil';
rate(3).iter_ans = 'n';
rate(3).name = 'Vpdh';
rate(3).units = 'umol/min/g';
rate(3).notes = 'Notes';
rate(3).guess = 0.000000e+00;
rate(3).delta = 1.000000e+00;
rate(3).upper_limit = 1.000000e+07;
rate(3).lower_limit = 0.000000e+00;

rate(4).V = 1.458000e-05;
rate(4).Expression = 'Rdil*Vtca';
rate(4).iter_ans = 'n';
rate(4).name = 'Vdil';
rate(4).units = 'umol/min/g';
rate(4).notes = 'Notes';
rate(4).guess = 0.000000e+00;
rate(4).delta = 1.000000e+00;
rate(4).upper_limit = 1.000000e+07;
rate(4).lower_limit = 0.000000e+00;

rate(5).V = 2.000000e-05;
rate(5).Expression = '';
rate(5).iter_ans = 'y';
rate(5).name = 'Rdil';
rate(5).units = 'umol/min/g';
rate(5).notes = 'Notes';
rate(5).guess = 0.000000e+00;
rate(5).delta = 1.000000e+00;
rate(5).upper_limit = 9.000000e-01;
rate(5).lower_limit = 0.000000e+00;

rate(6).V = 2.645000e-01;
rate(6).Expression = 'Vtca/2-0.1';
rate(6).iter_ans = 'n';
```

```
rate(6).name = 'Vgln';
rate(6).units = 'umol/min/g';
rate(6).notes = 'Notes';
rate(6).guess = 0.000000e+00;
rate(6).delta = 3.000000e-01;
rate(6).upper_limit = 1.000000e+00;
rate(6).lower_limit = 0.000000e+00;

rate(7).V = 1.500000e+00;
rate(7).Expression = '';
rate(7).iter_ans = 'y';
rate(7).name = 'Vlacin';
rate(7).units = 'umol/min/g';
rate(7).notes = 'Notes';
rate(7).guess = 0.000000e+00;
rate(7).delta = 1.000000e+00;
rate(7).upper_limit = 1.000000e+07;
rate(7).lower_limit = 0.000000e+00;

rate(8).V = 1.500000e+00;
rate(8).Expression = 'Vlacin';
rate(8).iter_ans = 'n';
rate(8).name = 'Vlacout';
rate(8).units = 'umol/min/g';
rate(8).notes = 'Notes';
rate(8).guess = 0.000000e+00;
rate(8).delta = 4.950000e-01;
rate(8).upper_limit = 1.000000e+07;
rate(8).lower_limit = 0.000000e+00;
%/**** Pools ****************************************/
conc(1).duplicate = 0;
conc(1).Conc = 6.700000e-01;
conc(1).Expression = '';
	conc(1).inflows(1).source_index = 1;
	conc(1).inflows(1).source_type = 1;
	conc(1).inflows(1).flow_multiplier = 2.000000;
	conc(1).inflows(1).number_isotopomers = 0;
	conc(1).inflows(1).match_index = -1;
	conc(1).inflows(1).MakeOnlyOne_flag = 0;
	conc(1).inflows(1).isot_fraction = 0.500;
	conc(1).inflows(1).rate_index = 1;
	conc(1).inflows(1).no_extra_mass = 0;
	conc(1).inflows(1).fate_map = [];
	conc(1).inflows(2).source_index = 2;
	conc(1).inflows(2).source_type = 1;
	conc(1).inflows(2).flow_multiplier = 1.000000;
	conc(1).inflows(2).number_isotopomers = 0;
	conc(1).inflows(2).match_index = -1;
	conc(1).inflows(2).MakeOnlyOne_flag = 0;
	conc(1).inflows(2).isot_fraction = 1.000;
	conc(1).inflows(2).rate_index = 7;
	conc(1).inflows(2).no_extra_mass = 0;
	conc(1).inflows(2).fate_map = [];
```

```
		conc(1).outflows(1).rate_index = 8;
		conc(1).outflows(2).rate_index = 3;
	conc(1).iter_ans = 'n';
	conc(1).number_carbons = 0;
	conc(1).carbon_position = '3,3';
	conc(1).MBVerified = 0;
	conc(1).isotopomer = 0;
	conc(1).start_APE = 0.000000;
	conc(1).T1 = 0.000000;
	conc(1).name = 'Lac';
	conc(1).units = 'umol/g';
	conc(1).notes = 'Concentration from QELT measurement in umol/g';
	conc(1).guess = 0.000000;
	conc(1).delta = 0.000000;
	conc(1).upper_limit = 10000000.000000;
	conc(1).lower_limit = 0.000000;
	conc(1).isotopomer = 0;

	conc(2).duplicate = 0;
	conc(2).Conc = 8.800000e+00;
	conc(2).Expression = '';
		conc(2).inflows(1).source_index = 1;
		conc(2).inflows(1).source_type = 0;
		conc(2).inflows(1).flow_multiplier = 1.000000;
		conc(2).inflows(1).number_isotopomers = 0;
		conc(2).inflows(1).match_index = -1;
		conc(2).inflows(1).MakeOnlyOne_flag = 0;
		conc(2).inflows(1).isot_fraction = 1.000;
		conc(2).inflows(1).rate_index = 3;
		conc(2).inflows(1).no_extra_mass = 0;
		conc(2).inflows(1).fate_map = [];
		conc(2).inflows(2).source_index = 2;
		conc(2).inflows(2).source_type = 1;
		conc(2).inflows(2).flow_multiplier = 1.000000;
		conc(2).inflows(2).number_isotopomers = 0;
		conc(2).inflows(2).match_index = -1;
		conc(2).inflows(2).MakeOnlyOne_flag = 0;
		conc(2).inflows(2).isot_fraction = 1.000;
		conc(2).inflows(2).rate_index = 4;
		conc(2).inflows(2).no_extra_mass = 0;
		conc(2).inflows(2).fate_map = [];
		conc(2).inflows(3).source_index = 3;
		conc(2).inflows(3).source_type = 0;
		conc(2).inflows(3).flow_multiplier = 1.000000;
		conc(2).inflows(3).number_isotopomers = 0;
		conc(2).inflows(3).match_index = -1;
		conc(2).inflows(3).MakeOnlyOne_flag = 0;
		conc(2).inflows(3).isot_fraction = 1.000;
		conc(2).inflows(3).rate_index = 6;
		conc(2).inflows(3).no_extra_mass = 0;
		conc(2).inflows(3).fate_map = [];
		conc(2).outflows(1).rate_index = 6;
		conc(2).outflows(2).rate_index = 2;
```

```
conc(2).iter_ans = 'n';
conc(2).number_carbons = 0;
conc(2).carbon_position = '4';
conc(2).MBVerified = 0;
conc(2).isotopomer = 0;
conc(2).start_APE = 0.000000;
conc(2).T1 = 0.000000;
conc(2).name = 'Glu';
conc(2).units = 'umol/g';
conc(2).notes = 'Concentration from QELT measurement in umol/g';
conc(2).guess = 0.000000;
conc(2).delta = 0.000000;
conc(2).upper_limit = 10000000.000000;
conc(2).lower_limit = 0.000000;
conc(2).isotopomer = 0;

conc(3).duplicate = 0;
conc(3).Conc = 5.820000e+00;
conc(3).Expression = '';
        conc(3).inflows(1).source_index = 2;
        conc(3).inflows(1).source_type = 0;
        conc(3).inflows(1).flow_multiplier = 1.000000;
        conc(3).inflows(1).number_isotopomers = 0;
        conc(3).inflows(1).match_index = -1;
        conc(3).inflows(1).MakeOnlyOne_flag = 0;
        conc(3).inflows(1).isot_fraction = 1.000;
        conc(3).inflows(1).rate_index = 6;
        conc(3).inflows(1).no_extra_mass = 0;
        conc(3).inflows(1).fate_map = [];
        conc(3).outflows(1).rate_index = 6;
conc(3).iter_ans = 'n';
conc(3).number_carbons = 0;
conc(3).carbon_position = '4';
conc(3).MBVerified = 0;
conc(3).isotopomer = 0;
conc(3).start_APE = 0.000000;
conc(3).T1 = 0.000000;
conc(3).name = 'Gln';
conc(3).units = 'umol/g';
conc(3).notes = 'Concentration from QELT measurement in umol/g';
conc(3).guess = 0.000000;
conc(3).delta = 0.000000;
conc(3).upper_limit = 10000000.000000;
conc(3).lower_limit = 0.000000;
conc(3).isotopomer = 0;
%/**** Combinations*************************************/
comb(1).name = 'Glx';
comb(1).Comb = 14.620000;
comb(1).units = 'umol/g';
comb(1).notes = 'Concentrations measured with QELT';
comb(1).number_pools = 2;
        comb(1).pools(1).pool_index = 2;
        comb(1).pools(1).type = 0;
```

```
		comb(1).pools(1).main_flag = 1;
		comb(1).pools(1).multiplier = 1.000000;
		comb(1).pools(2).pool_index = 3;
		comb(1).pools(2).type = 0;
		comb(1).pools(2).main_flag = 1;
		comb(1).pools(2).multiplier = 1.000000;
%/**** Drives ****************************************/
driv(1).name = 'Glc';
driv(1).units = 'mM';
driv(1).notes = 'Concentrations measured with DMI';
driv(1).number_carbons = 0;
driv(1).carbon_position = '6,6';
driv(1).T1 = 0.000000e+00;
driv(1).mirror_flag = 0;
		driv(1).outflows(1).rate_index = 1;
driv(1).x(1) = 0.000000e+00;	driv(1).y(1) = 0.000000e+00;	driv(1).z(1) = 0.000000e+00;
driv(1).x(2) = 1.000000e+01;	driv(1).y(2) = 8.256000e-01;	driv(1).z(2) = 7.191000e+01;
driv(1).x(3) = 2.000000e+01;	driv(1).y(3) = 1.105000e+00;	driv(1).z(3) = 9.624000e+01;
driv(1).x(4) = 3.000000e+01;	driv(1).y(4) = 1.464500e+00;	driv(1).z(4) = 1.275500e+02;
driv(1).x(5) = 4.000000e+01;	driv(1).y(5) = 1.618500e+00;	driv(1).z(5) = 1.409600e+02;
driv(1).x(6) = 5.000000e+01;	driv(1).y(6) = 1.795800e+00;	driv(1).z(6) = 1.564100e+02;
driv(1).x(7) = 6.000000e+01;	driv(1).y(7) = 1.873700e+00;	driv(1).z(7) = 1.631900e+02;
driv(1).x(8) = 7.000000e+01;	driv(1).y(8) = 1.866600e+00;	driv(1).z(8) = 1.625700e+02;
driv(1).x(9) = 8.000000e+01;	driv(1).y(9) = 1.869700e+00;	driv(1).z(9) = 1.628400e+02;
driv(1).x(10) = 9.000000e+01;	driv(1).y(10) = 1.876200e+00;	driv(1).z(10) = 1.634100e+02;

driv(2).name = 'NA';
driv(2).units = 'mM';
driv(2).notes = 'Notes';
driv(2).number_carbons = 0;
driv(2).carbon_position = '0';
driv(2).T1 = 0.000000e+00;
driv(2).mirror_flag = 0;
		driv(2).outflows(1).rate_index = 4;
		driv(2).outflows(2).rate_index = 7;
driv(2).x(1) = 0.000000e+00;	driv(2).y(1) = 1.000000e+00;	driv(2).z(1) = 1.150000e-02;
driv(2).x(2) = 1.000000e+00;	driv(2).y(2) = 1.000000e+00;	driv(2).z(2) = 1.150000e-02;
driv(2).x(3) = 1.000000e+02;	driv(2).y(3) = 1.000000e+00;	driv(2).z(3) = 1.150000e-02;
%/**** Conditions ****************************************/
initcond.Tmax = 1.000000e+02;
initcond.start_enrichment = 0.000000e+00;
initcond.max_step_size = 1.000000e+01;
initcond.IntTolAbs = 1.000000e-03;
initcond.IntTolRel = 1.000000e-03;
initcond.maxiter = 2;
initcond.ParallelProcessing = 1;
initcond.ProcessorsAvailable = 4;
initcond.ProcessorsSelected = 4;
initcond.ParProcBatchSize = 100;
initcond.startguess_delta = 25.000000;
%/**** Targets ****************************************/
targ(1).name = 'Glx C#';
targ(1).convert_factor = 1.335000e+00;
```

```
targ(1).match_rate_index = [];
targ(1).upper_limit = 0.000000e+00;
targ(1).lower_limit = 0.000000e+00;
targ(1).iter_ans = 'n';
targ(1).fit_weight = 1.000000e+00;
targ(1).type = 0;
targ(1).conc_index = 0;
targ(1).isot_index = 0;
targ(1).comb_conc_index = 1;
targ(1).comb_isot_index = 1;
        targ(1).x(1) = 0.000000e+00;        targ(1).y(1) = 0.000000e+00;
        targ(1).x(2) = 1.000000e+01;        targ(1).y(2) = 4.028000e-01;
        targ(1).x(3) = 2.000000e+01;        targ(1).y(3) = 6.520000e-01;
        targ(1).x(4) = 3.000000e+01;        targ(1).y(4) = 9.464000e-01;
        targ(1).x(5) = 4.000000e+01;        targ(1).y(5) = 1.193200e+00;
        targ(1).x(6) = 5.000000e+01;        targ(1).y(6) = 1.528300e+00;
        targ(1).x(7) = 6.000000e+01;        targ(1).y(7) = 1.790700e+00;
        targ(1).x(8) = 7.000000e+01;        targ(1).y(8) = 1.973800e+00;
        targ(1).x(9) = 8.000000e+01;        targ(1).y(9) = 2.175400e+00;
        targ(1).x(10) = 9.000000e+01;       targ(1).y(10) = 2.386500e+00;

targ(2).name = 'Lac C3,3';
targ(2).convert_factor = 1.335000e+00;
targ(2).match_rate_index = [];
targ(2).upper_limit = 0.000000e+00;
targ(2).lower_limit = 0.000000e+00;
targ(2).iter_ans = 'n';
targ(2).fit_weight = 1.000000e+00;
targ(2).type = 0;
targ(2).conc_index = 1;
targ(2).isot_index = 1;
targ(2).comb_conc_index = 0;
targ(2).comb_isot_index = 0;
        targ(2).x(1) = 0.000000e+00;        targ(2).y(1) = 0.000000e+00;
        targ(2).x(2) = 1.000000e+01;        targ(2).y(2) = 2.790000e-02;
        targ(2).x(3) = 2.000000e+01;        targ(2).y(3) = 1.866000e-01;
        targ(2).x(4) = 3.000000e+01;        targ(2).y(4) = 2.815000e-01;
        targ(2).x(5) = 4.000000e+01;        targ(2).y(5) = 3.857000e-01;
        targ(2).x(6) = 5.000000e+01;        targ(2).y(6) = 5.149000e-01;
        targ(2).x(7) = 6.000000e+01;        targ(2).y(7) = 5.771000e-01;
        targ(2).x(8) = 7.000000e+01;        targ(2).y(8) = 6.841000e-01;
        targ(2).x(9) = 8.000000e+01;        targ(2).y(9) = 7.650000e-01;
        targ(2).x(10) = 9.000000e+01;       targ(2).y(10) = 9.003000e-01;
%/**** Notes ***************************************/
notes = [];
```

*Model 2*

```
function [comb,conc,driv,rate,initcond,targ,notes] = fun(x)

%/**** Rates ****************************************/
```

```
rate(1).V = 3.645000e-01;
rate(1).Expression = 'Vtca/2';
rate(1).iter_ans = 'n';
rate(1).name = 'CMR';
rate(1).units = 'umol/min/g';
rate(1).notes = 'Notes';
rate(1).guess = 0.000000e+00;
rate(1).delta = 1.000000e+00;
rate(1).upper_limit = 1.000000e+07;
rate(1).lower_limit = 0.000000e+00;

rate(2).V = 7.290000e-01;
rate(2).Expression = '';
rate(2).iter_ans = 'y';
rate(2).name = 'Vtca';
rate(2).units = 'umol/min/g';
rate(2).notes = 'Mason et al. (1995)';
rate(2).guess = 0.000000e+00;
rate(2).delta = 3.000000e-01;
rate(2).upper_limit = 2.000000e+00;
rate(2).lower_limit = 0.000000e+00;

rate(3).V = 7.289854e-01;
rate(3).Expression = 'Vtca-Vdil';
rate(3).iter_ans = 'n';
rate(3).name = 'Vpdh';
rate(3).units = 'umol/min/g';
rate(3).notes = 'Notes';
rate(3).guess = 0.000000e+00;
rate(3).delta = 1.000000e+00;
rate(3).upper_limit = 1.000000e+07;
rate(3).lower_limit = 0.000000e+00;

rate(4).V = 1.458000e-05;
rate(4).Expression = 'Rdil*Vtca';
rate(4).iter_ans = 'n';
rate(4).name = 'Vdil';
rate(4).units = 'umol/min/g';
rate(4).notes = 'Notes';
rate(4).guess = 0.000000e+00;
rate(4).delta = 1.000000e+00;
rate(4).upper_limit = 1.000000e+07;
rate(4).lower_limit = 0.000000e+00;

rate(5).V = 2.000000e-05;
rate(5).Expression = '';
rate(5).iter_ans = 'y';
rate(5).name = 'Rdil';
rate(5).units = 'umol/min/g';
rate(5).notes = 'Notes';
rate(5).guess = 0.000000e+00;
rate(5).delta = 1.000000e+00;
rate(5).upper_limit = 9.000000e-01;
```

```
rate(5).lower_limit = 0.000000e+00;

rate(6).V = 4.660000e-01;
rate(6).Expression = '';
rate(6).iter_ans = 'y';
rate(6).name = 'Vgln';
rate(6).units = 'umol/min/g';
rate(6).notes = 'Mason et al. (1995)';
rate(6).guess = 0.000000e+00;
rate(6).delta = 3.000000e-01;
rate(6).upper_limit = 1.000000e+00;
rate(6).lower_limit = 0.000000e+00;

rate(7).V = 1.500000e+00;
rate(7).Expression = '';
rate(7).iter_ans = 'y';
rate(7).name = 'Vlacin';
rate(7).units = 'umol/min/g';
rate(7).notes = 'Notes';
rate(7).guess = 0.000000e+00;
rate(7).delta = 1.000000e+00;
rate(7).upper_limit = 1.000000e+07;
rate(7).lower_limit = 0.000000e+00;

rate(8).V = 1.500000e+00;
rate(8).Expression = 'Vlacin';
rate(8).iter_ans = 'n';
rate(8).name = 'Vlacout';
rate(8).units = 'umol/min/g';
rate(8).notes = 'Notes';
rate(8).guess = 0.000000e+00;
rate(8).delta = 4.950000e-01;
rate(8).upper_limit = 1.000000e+07;
rate(8).lower_limit = 0.000000e+00;
%/**** Pools ****************************************/
conc(1).duplicate = 0;
conc(1).Conc = 6.700000e-01;
conc(1).Expression = '';
        conc(1).inflows(1).source_index = 1;
        conc(1).inflows(1).source_type = 1;
        conc(1).inflows(1).flow_multiplier = 2.000000;
        conc(1).inflows(1).number_isotopomers = 0;
        conc(1).inflows(1).match_index = -1;
        conc(1).inflows(1).MakeOnlyOne_flag = 0;
        conc(1).inflows(1).isot_fraction = 0.500;
        conc(1).inflows(1).rate_index = 1;
        conc(1).inflows(1).no_extra_mass = 0;
        conc(1).inflows(1).fate_map = [];
        conc(1).inflows(2).source_index = 2;
        conc(1).inflows(2).source_type = 1;
        conc(1).inflows(2).flow_multiplier = 1.000000;
        conc(1).inflows(2).number_isotopomers = 0;
        conc(1).inflows(2).match_index = -1;
```

```
        conc(1).inflows(2).MakeOnlyOne_flag = 0;
        conc(1).inflows(2).isot_fraction = 1.000;
        conc(1).inflows(2).rate_index = 7;
        conc(1).inflows(2).no_extra_mass = 0;
        conc(1).inflows(2).fate_map = [];
        conc(1).outflows(1).rate_index = 8;
        conc(1).outflows(2).rate_index = 3;
conc(1).iter_ans = 'n';
conc(1).number_carbons = 0;
conc(1).carbon_position = '3,3';
conc(1).MBVerified = 0;
conc(1).isotopomer = 0;
conc(1).start_APE = 0.000000;
conc(1).T1 = 0.000000;
conc(1).name = 'Lac';
conc(1).units = 'umol/g';
conc(1).notes = 'Concentration from QELT measurement in umol/g';
conc(1).guess = 0.000000;
conc(1).delta = 0.000000;
conc(1).upper_limit = 10000000.000000;
conc(1).lower_limit = 0.000000;
conc(1).isotopomer = 0;

conc(2).duplicate = 0;
conc(2).Conc = 8.800000e+00;
conc(2).Expression = '';
        conc(2).inflows(1).source_index = 1;
        conc(2).inflows(1).source_type = 0;
        conc(2).inflows(1).flow_multiplier = 1.000000;
        conc(2).inflows(1).number_isotopomers = 0;
        conc(2).inflows(1).match_index = -1;
        conc(2).inflows(1).MakeOnlyOne_flag = 0;
        conc(2).inflows(1).isot_fraction = 1.000;
        conc(2).inflows(1).rate_index = 3;
        conc(2).inflows(1).no_extra_mass = 0;
        conc(2).inflows(1).fate_map = [];
        conc(2).inflows(2).source_index = 2;
        conc(2).inflows(2).source_type = 1;
        conc(2).inflows(2).flow_multiplier = 1.000000;
        conc(2).inflows(2).number_isotopomers = 0;
        conc(2).inflows(2).match_index = -1;
        conc(2).inflows(2).MakeOnlyOne_flag = 0;
        conc(2).inflows(2).isot_fraction = 1.000;
        conc(2).inflows(2).rate_index = 4;
        conc(2).inflows(2).no_extra_mass = 0;
        conc(2).inflows(2).fate_map = [];
        conc(2).inflows(3).source_index = 3;
        conc(2).inflows(3).source_type = 0;
        conc(2).inflows(3).flow_multiplier = 1.000000;
        conc(2).inflows(3).number_isotopomers = 0;
        conc(2).inflows(3).match_index = -1;
        conc(2).inflows(3).MakeOnlyOne_flag = 0;
        conc(2).inflows(3).isot_fraction = 1.000;
```

```
		conc(2).inflows(3).rate_index = 6;
		conc(2).inflows(3).no_extra_mass = 0;
		conc(2).inflows(3).fate_map = [];
		conc(2).outflows(1).rate_index = 6;
		conc(2).outflows(2).rate_index = 2;
conc(2).iter_ans = 'n';
conc(2).number_carbons = 0;
conc(2).carbon_position = '4';
conc(2).MBVerified = 0;
conc(2).isotopomer = 0;
conc(2).start_APE = 0.000000;
conc(2).T1 = 0.000000;
conc(2).name = 'Glu';
conc(2).units = 'umol/g';
conc(2).notes = 'Concentration from QELT measurement in umol/g';
conc(2).guess = 0.000000;
conc(2).delta = 0.000000;
conc(2).upper_limit = 10000000.000000;
conc(2).lower_limit = 0.000000;
conc(2).isotopomer = 0;

conc(3).duplicate = 0;
conc(3).Conc = 5.820000e+00;
conc(3).Expression = '';
		conc(3).inflows(1).source_index = 2;
		conc(3).inflows(1).source_type = 0;
		conc(3).inflows(1).flow_multiplier = 1.000000;
		conc(3).inflows(1).number_isotopomers = 0;
		conc(3).inflows(1).match_index = -1;
		conc(3).inflows(1).MakeOnlyOne_flag = 0;
		conc(3).inflows(1).isot_fraction = 1.000;
		conc(3).inflows(1).rate_index = 6;
		conc(3).inflows(1).no_extra_mass = 0;
		conc(3).inflows(1).fate_map = [];
		conc(3).outflows(1).rate_index = 6;
conc(3).iter_ans = 'n';
conc(3).number_carbons = 0;
conc(3).carbon_position = '4';
conc(3).MBVerified = 0;
conc(3).isotopomer = 0;
conc(3).start_APE = 0.000000;
conc(3).T1 = 0.000000;
conc(3).name = 'Gln';
conc(3).units = 'umol/g';
conc(3).notes = 'Concentration from QELT measurement in umol/g';
conc(3).guess = 0.000000;
conc(3).delta = 0.000000;
conc(3).upper_limit = 10000000.000000;
conc(3).lower_limit = 0.000000;
conc(3).isotopomer = 0;
%/**** Combinations*************************************/
comb(1).name = 'Glx';
comb(1).Comb = 14.620000;
```

```
comb(1).units = 'umol/g';
comb(1).notes = 'Concentrations measured with DMI';
comb(1).number_pools = 2;
        comb(1).pools(1).pool_index = 2;
        comb(1).pools(1).type = 0;
        comb(1).pools(1).main_flag = 1;
        comb(1).pools(1).multiplier = 1.000000;
        comb(1).pools(2).pool_index = 3;
        comb(1).pools(2).type = 0;
        comb(1).pools(2).main_flag = 1;
        comb(1).pools(2).multiplier = 1.000000;
%/**** Drives ****************************************/
driv(1).name = 'Glc';
driv(1).units = 'mM';
driv(1).notes = 'Concentrations measured with QELT';
driv(1).number_carbons = 0;
driv(1).carbon_position = '6,6';
driv(1).T1 = 0.000000e+00;
driv(1).mirror_flag = 0;
        driv(1).outflows(1).rate_index = 1;
driv(1).x(1) = 0.000000e+00;    driv(1).y(1) = 0.000000e+00;    driv(1).z(1) = 0.000000e+00;
driv(1).x(2) = 1.000000e+01;    driv(1).y(2) = 8.256000e-01;    driv(1).z(2) = 7.191000e+01;
driv(1).x(3) = 2.000000e+01;    driv(1).y(3) = 1.105000e+00;    driv(1).z(3) = 9.624000e+01;
driv(1).x(4) = 3.000000e+01;    driv(1).y(4) = 1.464500e+00;    driv(1).z(4) = 1.275500e+02;
driv(1).x(5) = 4.000000e+01;    driv(1).y(5) = 1.618500e+00;    driv(1).z(5) = 1.409600e+02;
driv(1).x(6) = 5.000000e+01;    driv(1).y(6) = 1.795800e+00;    driv(1).z(6) = 1.564100e+02;
driv(1).x(7) = 6.000000e+01;    driv(1).y(7) = 1.873700e+00;    driv(1).z(7) = 1.631900e+02;
driv(1).x(8) = 7.000000e+01;    driv(1).y(8) = 1.866600e+00;    driv(1).z(8) = 1.625700e+02;
driv(1).x(9) = 8.000000e+01;    driv(1).y(9) = 1.869700e+00;    driv(1).z(9) = 1.628400e+02;
driv(1).x(10) = 9.000000e+01;   driv(1).y(10) = 1.876200e+00;   driv(1).z(10) = 1.634100e+02;

driv(2).name = 'NA';
driv(2).units = 'mM';
driv(2).notes = 'Notes';
driv(2).number_carbons = 0;
driv(2).carbon_position = '0';
driv(2).T1 = 0.000000e+00;
driv(2).mirror_flag = 0;
        driv(2).outflows(1).rate_index = 4;
        driv(2).outflows(2).rate_index = 7;
driv(2).x(1) = 0.000000e+00;    driv(2).y(1) = 1.000000e+00;    driv(2).z(1) = 1.150000e-02;
driv(2).x(2) = 1.000000e+00;    driv(2).y(2) = 1.000000e+00;    driv(2).z(2) = 1.150000e-02;
driv(2).x(3) = 1.000000e+02;    driv(2).y(3) = 1.000000e+00;    driv(2).z(3) = 1.150000e-02;
%/**** Conditions ****************************************/
initcond.Tmax = 1.000000e+02;
initcond.start_enrichment = 0.000000e+00;
initcond.max_step_size = 1.000000e+01;
initcond.IntTolAbs = 1.000000e-03;
initcond.IntTolRel = 1.000000e-03;
initcond.maxiter = 2;
initcond.ParallelProcessing = 1;
initcond.ProcessorsAvailable = 4;
initcond.ProcessorsSelected = 4;
```

```
initcond.ParProcBatchSize = 100;
initcond.startguess_delta = 25.000000;
%/**** Targets ****************************************/
targ(1).name = 'Glx C#';
targ(1).convert_factor = 1.335000e+00;
targ(1).match_rate_index = [];
targ(1).upper_limit = 0.000000e+00;
targ(1).lower_limit = 0.000000e+00;
targ(1).iter_ans = 'n';
targ(1).fit_weight = 1.000000e+00;
targ(1).type = 0;
targ(1).conc_index = 0;
targ(1).isot_index = 0;
targ(1).comb_conc_index = 1;
targ(1).comb_isot_index = 1;
        targ(1).x(1) = 0.000000e+00;      targ(1).y(1) = 0.000000e+00;
        targ(1).x(2) = 1.000000e+01;      targ(1).y(2) = 4.028000e-01;
        targ(1).x(3) = 2.000000e+01;      targ(1).y(3) = 6.520000e-01;
        targ(1).x(4) = 3.000000e+01;      targ(1).y(4) = 9.464000e-01;
        targ(1).x(5) = 4.000000e+01;      targ(1).y(5) = 1.193200e+00;
        targ(1).x(6) = 5.000000e+01;      targ(1).y(6) = 1.528300e+00;
        targ(1).x(7) = 6.000000e+01;      targ(1).y(7) = 1.790700e+00;
        targ(1).x(8) = 7.000000e+01;      targ(1).y(8) = 1.973800e+00;
        targ(1).x(9) = 8.000000e+01;      targ(1).y(9) = 2.175400e+00;
        targ(1).x(10) = 9.000000e+01;     targ(1).y(10) = 2.386500e+00;

targ(2).name = 'Lac C3,3';
targ(2).convert_factor = 1.335000e+00;
targ(2).match_rate_index = [];
targ(2).upper_limit = 0.000000e+00;
targ(2).lower_limit = 0.000000e+00;
targ(2).iter_ans = 'n';
targ(2).fit_weight = 1.000000e+00;
targ(2).type = 0;
targ(2).conc_index = 1;
targ(2).isot_index = 1;
targ(2).comb_conc_index = 0;
targ(2).comb_isot_index = 0;
        targ(2).x(1) = 0.000000e+00;      targ(2).y(1) = 0.000000e+00;
        targ(2).x(2) = 1.000000e+01;      targ(2).y(2) = 2.790000e-02;
        targ(2).x(3) = 2.000000e+01;      targ(2).y(3) = 1.866000e-01;
        targ(2).x(4) = 3.000000e+01;      targ(2).y(4) = 2.815000e-01;
        targ(2).x(5) = 4.000000e+01;      targ(2).y(5) = 3.857000e-01;
        targ(2).x(6) = 5.000000e+01;      targ(2).y(6) = 5.149000e-01;
        targ(2).x(7) = 6.000000e+01;      targ(2).y(7) = 5.771000e-01;
        targ(2).x(8) = 7.000000e+01;      targ(2).y(8) = 6.841000e-01;
        targ(2).x(9) = 8.000000e+01;      targ(2).y(9) = 7.650000e-01;
        targ(2).x(10) = 9.000000e+01;     targ(2).y(10) = 9.003000e-01;

targ(3).name = 'Gln C4';
targ(3).convert_factor = 1.000000e+00;
targ(3).match_rate_index = [];
targ(3).upper_limit = 0.000000e+00;
```

```
targ(3).lower_limit = 0.000000e+00;
targ(3).iter_ans = 'n';
targ(3).fit_weight = 1.000000e+00;
targ(3).type = 1;
targ(3).conc_index = 3;
targ(3).isot_index = 3;
targ(3).comb_conc_index = 0;
targ(3).comb_isot_index = 0;
        targ(3).x(1) = 0.000000e+00;        targ(3).y(1) = 0.000000e+00;
        targ(3).x(2) = 5.000000e+00;        targ(3).y(2) = -4.400000e-01;
        targ(3).x(3) = 1.000000e+01;        targ(3).y(3) = 1.500000e-01;
        targ(3).x(4) = 1.500000e+01;        targ(3).y(4) = 2.510000e+00;
        targ(3).x(5) = 2.000000e+01;        targ(3).y(5) = 5.080000e+00;
        targ(3).x(6) = 2.500000e+01;        targ(3).y(6) = 4.020000e+00;
        targ(3).x(7) = 3.000000e+01;        targ(3).y(7) = 9.660000e+00;
        targ(3).x(8) = 3.500000e+01;        targ(3).y(8) = 1.327000e+01;
        targ(3).x(9) = 4.000000e+01;        targ(3).y(9) = 1.701000e+01;
        targ(3).x(10) = 4.500000e+01;       targ(3).y(10) = 1.753000e+01;
        targ(3).x(11) = 5.000000e+01;       targ(3).y(11) = 9.740000e+00;
        targ(3).x(12) = 5.500000e+01;       targ(3).y(12) = 1.078000e+01;
        targ(3).x(13) = 6.000000e+01;       targ(3).y(13) = 1.196000e+01;
        targ(3).x(14) = 6.500000e+01;       targ(3).y(14) = 1.654000e+01;
        targ(3).x(15) = 7.000000e+01;       targ(3).y(15) = 2.019000e+01;
        targ(3).x(16) = 7.500000e+01;       targ(3).y(16) = 1.956000e+01;
        targ(3).x(17) = 8.000000e+01;       targ(3).y(17) = 1.988000e+01;
        targ(3).x(18) = 8.500000e+01;       targ(3).y(18) = 2.132000e+01;

targ(4).name = 'Glu C4';
targ(4).convert_factor = 1.000000e+00;
targ(4).match_rate_index = [];
targ(4).upper_limit = 0.000000e+00;
targ(4).lower_limit = 0.000000e+00;
targ(4).iter_ans = 'n';
targ(4).fit_weight = 1.000000e+00;
targ(4).type = 1;
targ(4).conc_index = 2;
targ(4).isot_index = 2;
targ(4).comb_conc_index = 0;
targ(4).comb_isot_index = 0;
        targ(4).x(1) = 0.000000e+00;        targ(4).y(1) = 0.000000e+00;
        targ(4).x(2) = 5.000000e+00;        targ(4).y(2) = 9.300000e-01;
        targ(4).x(3) = 1.000000e+01;        targ(4).y(3) = 2.980000e+00;
        targ(4).x(4) = 1.500000e+01;        targ(4).y(4) = 5.590000e+00;
        targ(4).x(5) = 2.000000e+01;        targ(4).y(5) = 6.000000e+00;
        targ(4).x(6) = 2.500000e+01;        targ(4).y(6) = 9.490000e+00;
        targ(4).x(7) = 3.000000e+01;        targ(4).y(7) = 1.229000e+01;
        targ(4).x(8) = 3.500000e+01;        targ(4).y(8) = 1.264000e+01;
        targ(4).x(9) = 4.000000e+01;        targ(4).y(9) = 1.505000e+01;
        targ(4).x(10) = 4.500000e+01;       targ(4).y(10) = 1.805000e+01;
        targ(4).x(11) = 5.000000e+01;       targ(4).y(11) = 1.696000e+01;
        targ(4).x(12) = 5.500000e+01;       targ(4).y(12) = 1.827000e+01;
        targ(4).x(13) = 6.000000e+01;       targ(4).y(13) = 1.865000e+01;
        targ(4).x(14) = 6.500000e+01;       targ(4).y(14) = 2.188000e+01;
```

```
        targ(4).x(15) = 7.000000e+01;        targ(4).y(15) = 2.358000e+01;
        targ(4).x(16) = 7.500000e+01;        targ(4).y(16) = 2.442000e+01;
        targ(4).x(17) = 8.000000e+01;        targ(4).y(17) = 2.621000e+01;
        targ(4).x(18) = 8.500000e+01;        targ(4).y(18) = 2.795000e+01;
%/**** Notes ****************************************/
notes = [];
```

**Fig. S1:** Fitted time-curves using CWave

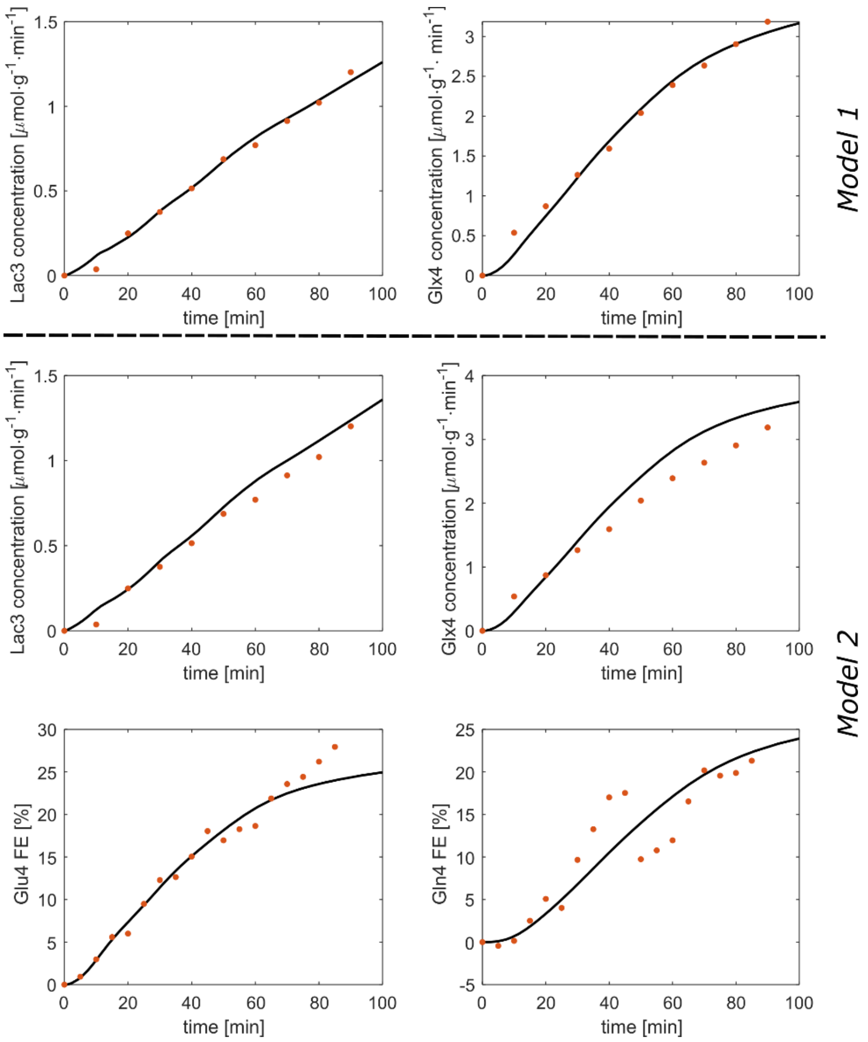


**Fig. S1:** Fitted time-curves using CWave. The first row shows the fitted time-curves as well as the measured data points for model 1. The lower rows show the corresponding time-curves for model 2.

**Fig. S2:** Glutamate and Glutamine VeSPA basis sets with and without $^2$H labeling

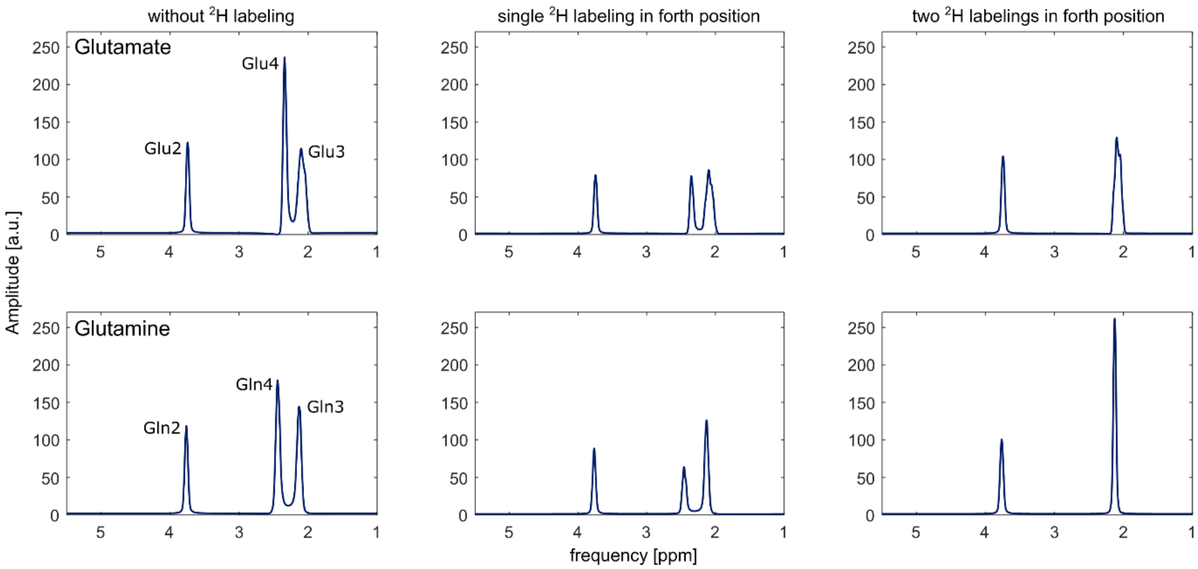


**Fig. S2:** Glutamate and Glutamine VeSPA basis sets with and without $^2$H labeling. The upper row shows the simulated $^1$H basis set for Glu with a varying number of $^2$H labels. The coupling between proton and deuterium was neglected (coupling constant 0 Hz). The lower row shows the corresponding basis sets for Gln.
